# Neural mechanisms to exploit positional geometry for collision avoidance

**DOI:** 10.1101/2021.12.11.472218

**Authors:** Ryosuke Tanaka, Damon A. Clark

## Abstract

Visual motion provides rich geometrical cues about the three-dimensional configuration the world. However, how brains decode the spatial information carried by motion signals remains poorly understood. Here, we study a collision avoidance behavior in *Drosophila* as a simple model of motion-based spatial vision. With simulations and psychophysics, we demonstrate that walking *Drosophila* exhibit a pattern of slowing to avoid collisions by exploiting the geometry of positional changes of objects on near-collision courses. This behavior requires the visual neuron LPLC1, whose tuning mirrors the behavior and whose activity drives slowing. LPLC1 pools inputs from object- and motion-detectors, and spatially biased inhibition tunes it to the geometry of collisions. Connectomic analyses identified circuitry downstream of LPLC1 that faithfully inherits its response properties. Overall, our results reveal how a small neural circuit solves a specific spatial vision task by combining distinct visual features to exploit universal geometrical constraints of the visual world.

## Introduction

The problem of spatial vision addresses how we can sense three-dimensional configurations of our surroundings from the “flat” images on our retinas. This problem has long been a central issue in vision science (Berkeley, 1709; von Helmholtz, 1924). In solving this problem, visual motion is a particularly useful source of spatial information, since the pattern of retinal motion caused by relative movements between an observer and its environment follows lawful geometry (Gibson, 1966). Indeed, neuroanatomical and physiological studies in primates have established that motion-sensitive cortical visual areas, like area MT, comprise a part of the so-called “where” pathway (Goodale and Milner, 1992; Mishkin et al., 1983) and contribute to the perception of three-dimensional structures based on motion cues (Bradley et al., 1998). However, circuit-level understanding of how spatial information carried by visual motion is decoded to guide specific behaviors remains largely missing. A useful model system to explore the mechanism of motion-based spatial vision is the fruit fly *Drosophila*, where powerful genetic (Guo et al., 2019) and connectomic (Scheffer et al., 2020) tools allow one to dissect neural circuit mechanisms in detail. In addition, recent years have seen rapid progress in understanding of the motion detection circuitry in the *Drosophila* visual system (Borst et al., 2020; Yang and Clandinin, 2018), which can now guide attempts to pinpoint neural mechanisms of spatial vision in flies.

For many animals, one routine task that requires spatial vision is avoiding collisions with other animals. Collision with predators poses an obvious survival threat to animals, and unwanted collisions with conspecifics compromise navigation, even when there is no risk of predation. As objects move relative to the observer, geometry dictates the size and position of their retinal images over time. Objects approaching the observer expand in apparent size, or ‘loom’, providing a useful and well-studied collision cue (Branco and Redgrave, 2020; Peek and Card, 2016). Importantly, beyond the change in size, change in an object’s position can also provide useful cues about impending collisions: provided an observer and an approaching object both maintain constant velocities, the retinal position of the object stays constant only if it is on a collision course, a situation analogous to “constant bearing, decreasing range” in maritime navigation (Murtaugh and Criel, 1966). Similarly, approaching objects will move back-to-front across the retina if they will cross in front of the observer, or will move front-to-back across the retina if they will cross behind the observer (**Fig. 1A**). Path crossings in front pose a collision risk to the observer, especially when the object crossing a path is capable of stopping suddenly. Thus, back-to-front motion can function as a heuristic geometrical cue for imminent future collisions. Indeed, a previous study demonstrated that walking flies halt upon observing visual objects moving back-to-front (Zabala et al., 2012), a strategy that could avoid collisions with conspecifics (Chalupka et al., 2016). However, the circuits governing such collision avoidance based on directional motion remain unknown.

**Figure 1.**
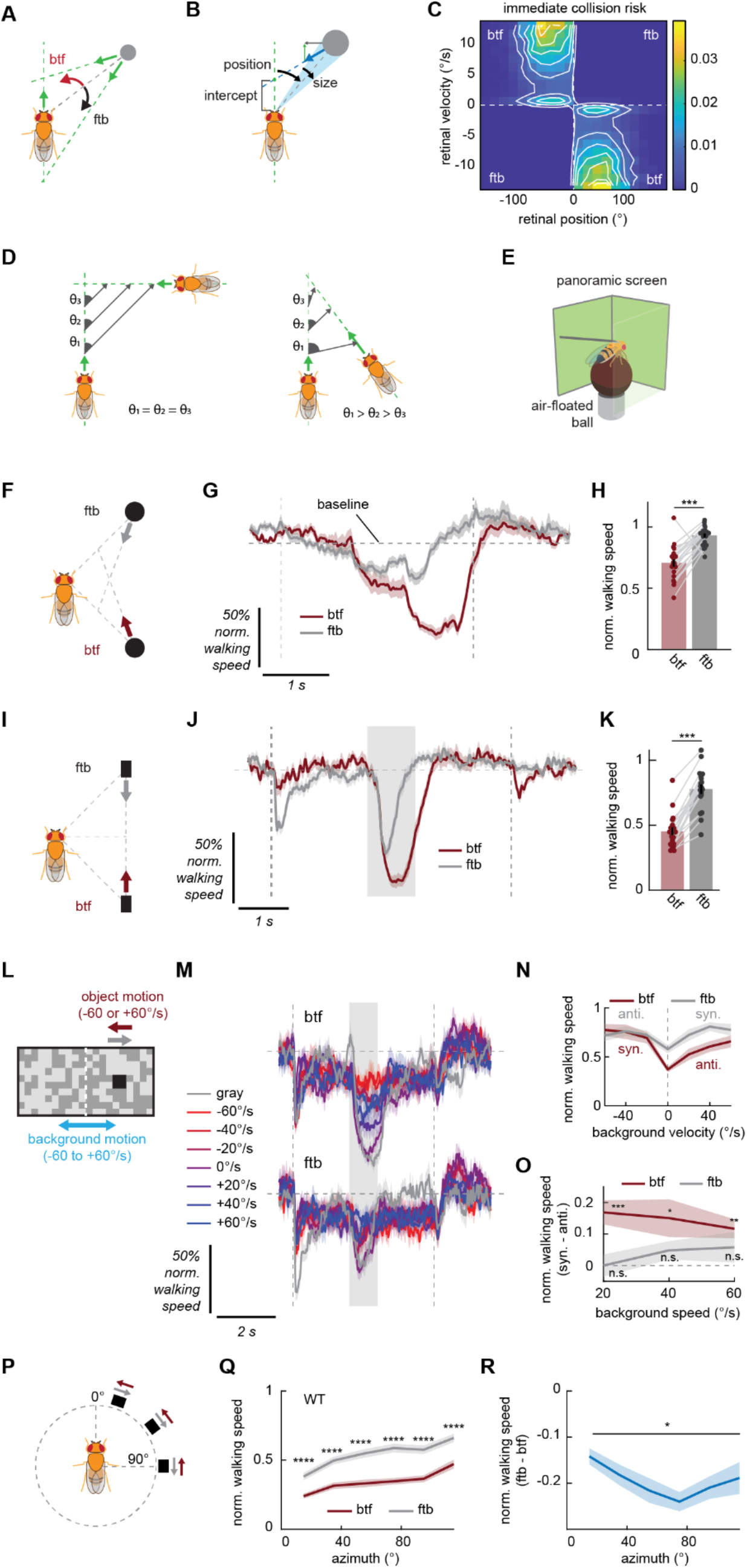
Flies exhibit slowing that mirrors geometry of collisions. (A) Geometry of collisions. Objects crossing the path in front of an observer appear to move in the back-to-front (btf) direction across the retina, whereas ones crossing behind the observer will appear to move front-to-back (ftb). (B) A schematic of the simulation. Linearly translating circular objects were placed at random around an observer that moved forward at a constant velocity. The collision risk posed by the object was calculated based on their future path-crossing intercept. (C) Immediate collision risk, defined as time-discounted inverse of positive future intercept (see *Methods* for details), as a function of angular position and velocity. Odd and even quadrants respectively correspond to front-to-back and back-to-front motion. (D) (*left*) When an object is on an exact collision course with the observer, the relative bearing (θ) of the observer remains constant. (*right*) An object that crosses the path in front of the observer at an acute angle decrease its bearing as they approach, causing back-to-front motion. (E) Schematic of the setup for the behavioral experiments in which flies walked on a spherical treadmill while they were presented with panoramic visual stimuli. (F) In the approach stimuli, simulated black circular objects approached the fly obliquely either from the front (ftb) or from the back (btf). (G, H) Wildtype fly normalized walking response to the approach stimuli in either direction, (G) as a function of time or (H) time-averaged. Forward walking speed was normalized by the baseline speed during the preceding interstimulus period, which is indicated by horizontal dotted line. The vertical dotted lines mark the beginning and the end of the stimulus. Each dot in (H) represents a fly, and data from the same flies are connected with gray lines. (I) In the parallel stimuli, simulated black rectangular objects appeared by the fly and remained stationary for 2 seconds, moved in a trajectory parallel to the fly in either direction for one second, stopped for another 2 seconds, and then disappeared. (J, K) Same as (G, H), but for the parallel stimuli. Time-averaged responses were calculated within the shaded region in (J). The vertical dotted lines and the shaded regions respectively represent on- and offset of the object and the period during which the object was moving. (L) Schematic of the stimuli used to test the interaction between the collision avoidance behavior and background motion. (M) Wildtype fly normalized walking response to squares moving in either direction (*top*: back-to-front, *bottom*: front-to-back), paired with rotating backgrounds. The velocity of the background is color-coded. The gray shaded region indicates when the object was moving. (N) Time-averaged normalized walking responses of wildtype flies to squares moving in either direction, as functions of background velocities. Averaging was within the shaded region in (M). Positive velocity is in the same direction as back-to-front (btf). (O) Time-averaged normalized walking speed in response to squares when the background was moving with the square minus when the background was moving against the square, for each background speed. (P) To probe retinotopic bias in the direction selective slowing, black rectangular objects sweeping short horizontal trajectories in either direction were presented at various azimuthal locations. (Q, R) Time-averaged normalized walking response of wildtype flies to the azimuth sweep stimuli as functions of azimuth, either (Q) by the motion directions or (R) the difference between the two. The averaging window was 1 second long from the onset of the stimuli. Error bars and shades around mean traces all indicate standard error of the mean. (G, H) N = 21 flies. (J, K) N = 19 flies. (M-O) N = 19 flies. (Q, R) N = 39 flies. n. s.: not significant (p> .05); *: p < .05; **: p < .01; ***: p < .001; ****: p < .0001 in Wilcoxon signed-rank test or Friedman test (R only).

Here, we investigate how *Drosophila* uses positional changes to avoid collision at both behavioral and circuit levels. First, by combining simulations and a high-throughput psychophysics, we demonstrate that the flies exhibit a pattern of slowing that avoids collisions by exploiting the positional geometry associated with them. Second, using synaptic silencing and optogenetics, we show that a visual projection neuron called LPLC1 is necessary for this collision avoidance behavior, and activating LPLC1 elicits slowing. LPLC1’s response properties, as measured with two-photon calcium imaging, mirror the tuning of the collision avoidance behavior, including a spatial bias in direction selectivity concordant with the positional geometry of collisions. Third, we show that LPLC1 combines excitatory inputs from elementary motion and object detectors, and achieves selectivity for objects on near-collision courses in part through spatially biased glutamatergic inhibition. Last, we identify a central brain pathway for this collision avoidance and show that it faithfully inherits response properties of LPLC1. Overall, the results reveal how signals from motion and object detectors can be combined to implement a solution for a spatial vision task that exploits a universal geometrical constraint of the visual world.

## Results

### Back-to-front motion is a useful terrestrial collision cue

As objects move relative to an observer, their apparent size and position change systematically as dictated by geometry. There are at least two reasons to think that back-to-front motion in particular can be a useful heuristic cue to detect and avoid collisions with objects. First, optic flow caused by forward translation always moves front-to-back. Therefore, any back-to-front motion observed during forward locomotion can be attributed to non-stationary objects in the surroundings (Zabala et al., 2012). Second, an approaching object will appear moving in the back-to-front direction if it is about to cross in front of the observer, and will appear to move front-to-back if it will cross behind the observer (**Fig. 1A**). If the approaching object is an animal, it could stop while crossing in front of the observer and thus poses a collision risk. To gain better intuition about how and when back-to-front motion is useful to predict frontal path crossings, we simulated an observer moving forward in the presence of objects with random relative positions and constant random velocities (**Fig. 1B**). We quantified how each object contributed to the ‘immediate collision risk’, defined as the time-discounted, inverted intercept between the observer and object trajectories (see *Methods* for details of the simulation).

When we plotted the collision risk against retinal angular position and velocity of the object (**Fig. 1C**), there were two pairs of clusters with high collision risk: one around zero velocity and the other around large velocities in the back-to-front direction. The zero-velocity clusters correspond to the “constant bearing, decreasing range” situation (**Fig. 1D**), where the object is directly intercepting the observer. A second cluster with higher back-to-front velocities represents nearby objects about to cross in front of the observer at acute angles (**Fig. 1D**). In these higher velocity clusters, the collision risk was higher for lateral rather than for directly frontal objects (**Fig. 1C**). This is because objects moving laterally right in front of the observer tend to cross the observer’s path long before the observer reaches that location. These results suggest that back-to-front motion of objects predicts imminent near-collisions, especially in the frontolateral visual field.

### Drosophila shows direction selective slowing in response to stimuli mimicking conspecifics

With the above geometrical results in mind, we designed experiments to characterize how flies respond to visual objects moving in the back-to-front direction in our high throughput psychophysics assay. In our assay, tethered flies were placed above of air-suspended balls, and their walking responses were recorded as the rotation of the balls (Creamer et al., 2019; Salazar-Gatzimas et al., 2016). Visual stimuli were presented on panoramic screens surrounding the flies (**Fig. 1E**). As visual stimuli, we first simulated a black object that linearly approached the fly from the side with a constant velocity, independent of the fly’s behavior (hereafter ‘approach stimulus’; See *Methods* for details). The size (2 mm tall, 3 mm wide) and velocity (20 mm/s) of the object was approximately matched to the realistic size and walking velocity of *Drosophila* (Branson et al., 2009; DeAngelis et al., 2019). The trajectories of the objects started either in front of or behind the fly in a symmetric manner, and only the objects starting from behind the fly are projected to cross in front of the observer fly. From the fly’s perspective, the objects appearing behind move back-to-front, while those appearing in front move front-to-back, and both expand in size identically over time (**Fig. 1F**). Wildtype flies slowed slightly in response to the front-starting approach stimulus, but slowed substantially more and for a longer duration in response to the rear-starting approach stimuli (**Fig. 1G, H**).

Although this observation is consistent with the idea that flies freeze in response to back-to-front moving objects, as previously observed (Zabala et al., 2012), it is also possible that the looming in the frontal visual field, rather than back-to-front motion itself, triggered the observed slowing. To exclude this possibility, we simulated a rectangular object that moved parallel to the fly, again starting either from in front of or behind the fly (hereafter ‘parallel stimulus’) (**Fig. 1I**). Wildtype flies presented with the parallel stimuli again slowed significantly more in response to rear-starting conditions (**Fig. 1J, K**). Since the front- and rear-starting parallel stimuli are trajectory-matched and contain virtually no looming, this result strongly suggests that the observed slowing behavior is selective for the direction of object motion. An object moving parallel to the fly’s trajectory constitutes a false-positive case from the collision avoidance perspective, since such a parallel trajectory would never cross the path of the observer, but flies have been reported to freeze in response to such stimuli (Zabala et al., 2012). In addition to slowing, both approach and parallel stimuli also elicited mild turning against the direction and position of object motion (**Fig. S1A, B**) (Maimon et al., 2008; Tanaka and Clark, 2020). Overall, these results suggest that flies initiate slowing in response to back-to-front motion, likely reflecting a collision avoidance behavior.

### The pattern of direction selective slowing mirrors the geometry of collision

One of the reasons why back-to-front motion can be a useful collision cue is that it is directed counter to the optic flow from forward translation and thus can be unambiguously attributed to moving objects. This argument suggests that flies would exhibit the direction-selective collision avoidance behavior even in the presence of cluttered, moving backgrounds, as long as objects and backgrounds are moving against each other. To test this hypothesis, we presented wildtype flies with 10° x 10° black squares translating against half-contrast, 5° resolution random checkerboard patterns that rotated around the fly at several velocities (**Fig. 1L**). Rotational, rather than translational background was used, since translational optic flow presented in an open-loop manner by itself potently slows flies (Creamer et al., 2018), making it difficult to observe additional slowing induced by objects. Overall, rotating backgrounds, especially fast ones, suppressed the slowing caused by moving objects (**Fig. 1M, N**), in addition to causing mild slowing and strong turning (**Fig. S1C**). Interestingly, while slowing caused by a front-to-back object was suppressed equally by backgrounds moving either direction, flies slowed significantly more to an object moving in the back-to-front direction when it is on a background moving against rather than with the object (**Fig. 1O**). This result indicates that flies use relative motion between object and background, in addition to the directionality of object motion itself, to initiate slowing.

Last, we asked if there is any spatial bias in the observed slowing behavior that could match the geometry of frontal path crossing (**Fig. 1C**). To do so, we presented a small square sweeping a short horizontal trajectory in either direction at different azimuthal locations (**Fig. 1P**). Objects in front elicited more slowing in wildtype flies regardless of direction, and slowing was selective for back-to-front direction at all azimuths (**Fig. 1Q**). However, the directional difference in slowing showed a U-shaped pattern, indicating that slowing in response to frontolateral objects were more selective for back-to-front direction (**Fig. 1R**). This result implies that the direction selectivity of the slowing behavior is strongest in the same azimuthal range where back-to-front motion most strongly predicts future collision (**Fig. 1C**).

### LPLC1 activity is necessary and sufficient for the slowing behavior

We next worked to identify neural substrates for this collision avoidance behavior. Since the slowing is selective for the direction of object motion, we hypothesized that synaptic outputs of T4 and T5 neurons, the first direction selective cells in the fly visual system (Maisak et al., 2013), would be necessary for the behavior. When we silenced the synaptic output of T4 and T5 by introducing *shibire^ts^* (Kitamoto, 2001) to these cells, slowing in response to back-to-front parallel stimuli was significantly reduced compared to the genetic controls (**Fig. 2A, B**), while slowing in response to front-to-back stimuli was significantly increased, almost abolishing the direction selectivity in the behavior. Similarly, silencing T4 and T5 significantly reduced fly slowing in response to back-to-front approach stimuli (**Fig. S1D, E**). These results show that T4/T5 are required for the direction selective collision avoidance behavior.

**Figure 2.**
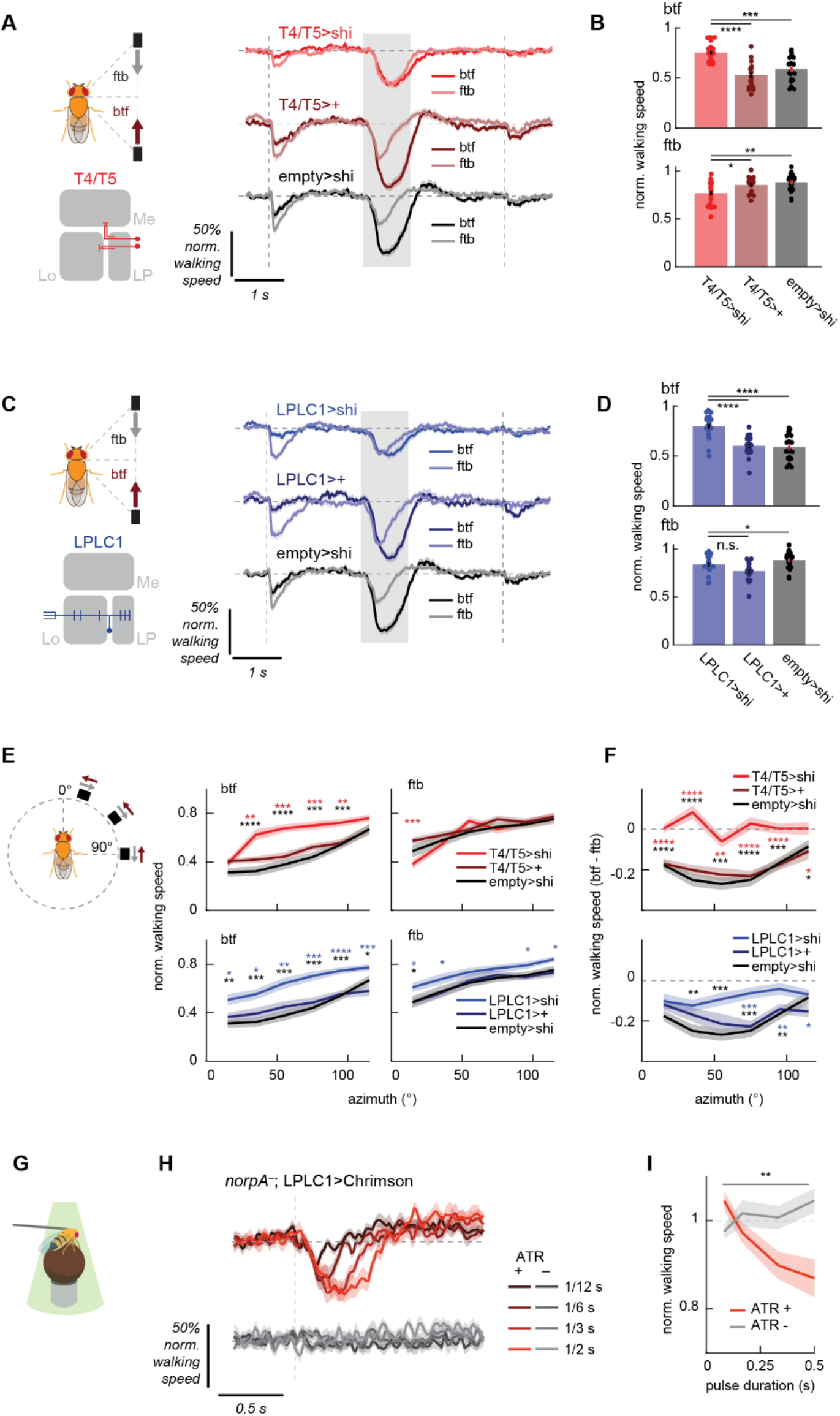
LPLC1 is necessary for collision avoidance and sufficient to cause slowing. (A, B) Normalized walking responses of T4/T5 silenced flies and their controls in response to the parallel stimuli, (A) over time or (B) averaged over time, as in Fig. 1J, K. (C, D) Same as (A, B), but for LPLC1. (E) Time-averaged walking responses of (*top*) T4/T5 or (*bottom*) LPLC1 silenced flies with their respective controls to the azimuth sweep stimuli by directions, as in Fig. 1Q. (F) The directional differences of the walking responses of the same flies as in (E) to the azimuth sweep stimuli, as in Fig. 1R. (G) A schematic of the optogenetics setup. (H, I) Walking response of LPLC1>Chrimson flies with or without ATR feeding to pulses of green light, either (H) over time or (I) time-averaged. The averaging window was 1 second long. Error bars and shades around mean traces all indicate standard error of the mean. (A, B) N = 19 (T4/T5>shi), 17 (T4/T5>+), 22 (empty>shi) flies. (C, D) N = 20 (LPLC1 >shi), 17 (LPLC1>+), 22 (empty>shi) flies. (E, F) N = 18 (T4/T5>shi), 20 (T4/T5>+), 19 (LPLC1>shi), 16 (LPLC1>+), 19 (empty>shi) flies. (H, I) N = 13 (ATR+), 12 (ATR-) flies. n. s.: not significant (p >.05); *: p < .05; **: p < .01; ***: p < .001; ****: p < .0001 in Wilcoxon rank sum test (B, D-F) and 2-way analysis of variance (ANOVA) (I; the main effect of ATR conditions).

Next, we aimed to identify neurons downstream of T4/T5 that selectively respond to objects moving back-to-front to trigger the slowing behavior. Lobula plate (LP), the neuropil where T4/T5 axon terminals reside, is innervated by several types of columnar visual projection neurons (VPNs) (Eliason, 2017; Fischbach and Dittrich, 1989; Isaacson, 2018; Mu et al., 2012; Panser et al., 2016; Wu et al., 2016). Columnar VPNs have been shown to detect specific local visual features that trigger a variety of behaviors (Ache et al., 2019; Eliason, 2017; Isaacson, 2018; Klapoetke et al., 2017; von Reyn et al., 2017; Ribeiro et al., 2018; Tanaka and Clark, 2020; Wu et al., 2016), so they make good candidates for the putative back-to-front moving object detector. Among the known LP-innervating columnar VPN types, LPLC2 has been shown to detect visual loom and drive escape responses (Ache et al., 2019; Klapoetke et al., 2017) and LPC1 and LLPC1 to detect translational optic flow and drive slowing (Eliason, 2017; Isaacson, 2018). Among remaining LP-innervating VPNs with no known function, a neuron type called lobula plate-lobula columnar cell type 1 (LPLC1) is particularly well positioned to detect objects moving back-to-front, because it innervates layer 2 of LP, which houses T4/T5 terminals tuned to back-to-front motion, but not the front-to-back-selective layer 1 (Maisak et al., 2013; Wu et al., 2016). To test whether LPLC1 is necessary for the slowing, we silenced synaptic outputs of LPLC1 by expressing *shibire^ts^*, and examined its effect on the behavior. We found that flies with LPLC1 silenced slowed significantly less in response to the back-to-front parallel (**Fig. 2C, D**) as well as approach stimuli (**Fig. S1F, G**), indicating that LPLC1 is necessary for the wild-type slowing phenotype. We also confirmed that silencing LPLC1 does not affect several visuomotor behaviors known to be dependent on T4/T5 (**Fig. S1H-J**).

We also tested how silencing either T4/T5 or LPLC1 affects the spatial bias in the slowing behavior (**Fig. 1P-R**). Silencing T4 and T5 increased slowing in response to front-to-back objects in front, and reduced slowing in response to front-to-back objects on the side (**Fig. 2E**). This reduced the direction selectivity of slowing across the almost all azimuth tested, abolishing the U-shaped pattern of directional difference in slowing visible in control genotypes (**Fig. 2F**). Similarly, silencing of LPLC1 reduced slowing in response to back-to-front objects across broad azimuths (**Fig. 2E**). However, reduction in directional differences of slowing was only significant from the both of the two control genotypes at lateral azimuths (**Fig. 2F**). This result suggests that direction selectivity of LPLC1 neurons is spatially biased and most pronounced in the frontolateral azimuthal range where back-to-front motion most strongly predicts near collision (**Fig. 1C**).

To further confirm the involvement of LPLC1 in the slowing behavior, we optogenetically activated LPLC1 neurons in blind (*norpA*^-^) flies, and tested whether activity in LPLC1 can trigger slowing. Blind flies expressing a red-shifted channelrhodopsin Chrimson (Klapoetke et al., 2014) in LPLC1 were tethered on air suspended balls, and pulses of green light with various durations were shone onto the flies from the DLP projectors (Creamer et al., 2019; Tanaka and Clark, 2020) (**Fig. 2G**). We compared the walking velocity changes in response to green lights between flies fed with food with or without all-trans retinal (ATR) (de Vries and Clandinin, 2013), a cofactor necessary for channelrhodopsin function. While flies fed with food without ATR did not show any response to green lights, flies fed with ATR exhibited duration-dependent slowing in response to green light (**Fig. 2H, I**), showing that the activity of LPLC1 alone is sufficient to make flies slow.

### Visual response properties of LPLC1 neurons mirror the tuning of the collision avoidance behavior

To better understand how LPLC1 contributes to this collision avoidance behavior, we next used two-photon calcium imaging to directly explore the visual tuning of LPLC1 neurons (**Fig. 3A**). First, to broadly characterize their response properties, we imaged the axon terminals of LPLC1 neurons expressing GCaMP6f (Chen et al., 2013) while presenting a variety of visual stimuli. The axon terminals of columnar VPNs including LPLC1 form structures called optic glomeruli, where retinotopy is mostly discarded (Otsuna and Ito, 2006; Panser et al., 2016; Wu et al., 2016 -- but see Morimoto et al., 2020). Thus, glomerular calcium activity can be interpreted as the spatially averaged population activity of LPLC1 neurons. We used a battery of stimuli consisting of full-field drifting square wave gratings, full-field flashes, moving bars and small squares, and expanding disks. LPLC1 did not respond to wide field stimuli, while it did respond to moving bars and small squares (**Fig. 3B**), consistent with previous measurements (Städele et al., 2020). As expected from the behavioral results, LPLC1’s responses to bars and squares were significantly selective for the back-to-front direction (**Fig. 3C**). LPLC1 vigorously responded to dark expanding disks, similar to several other types of columnar VPNs (Ache et al., 2019; Klapoetke et al., 2017; Morimoto et al., 2020; von Reyn et al., 2017; Wu et al., 2016).

**Figure 3.**
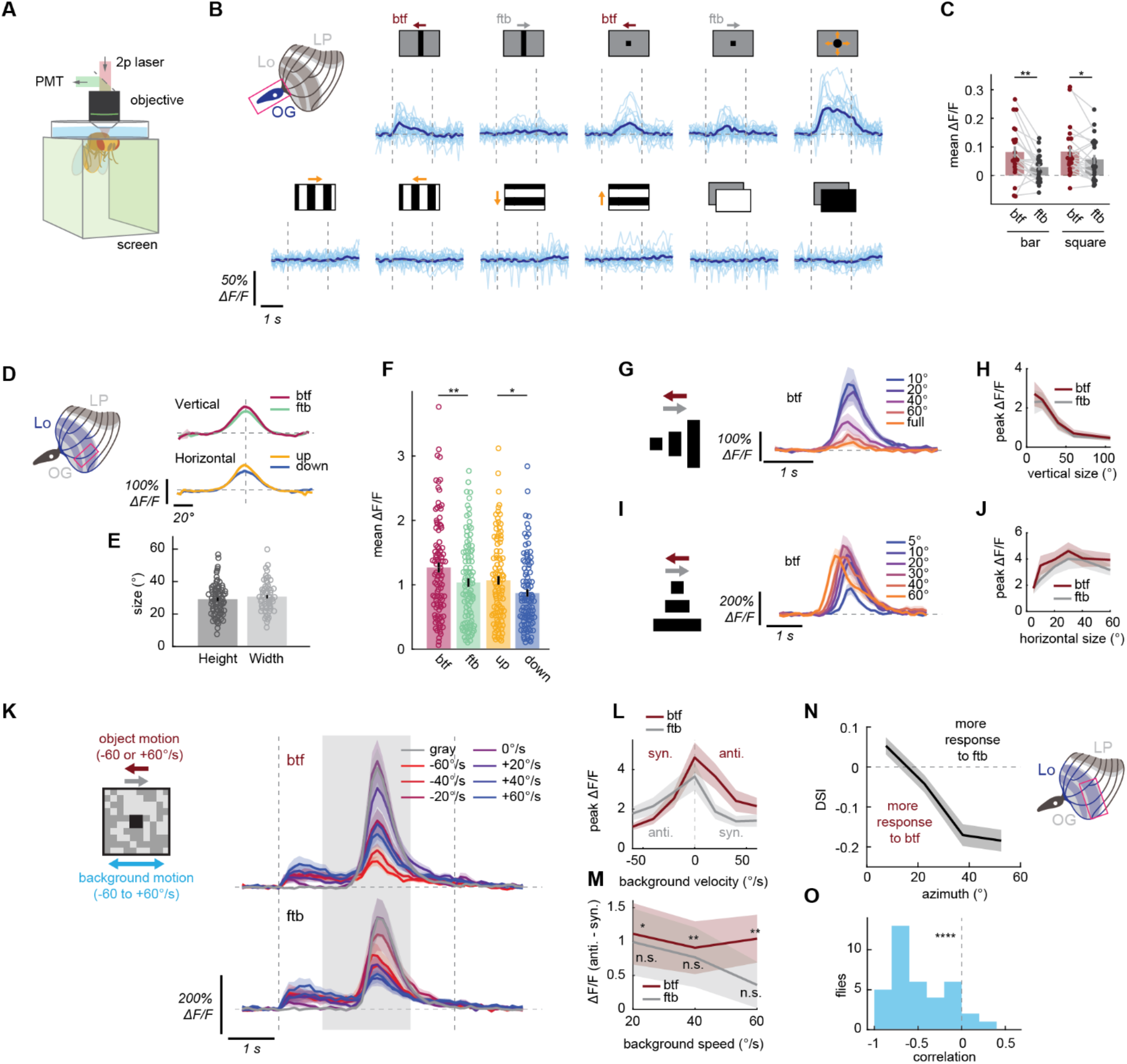
Physiological response properties of LPLC1 match the tuning of the collision avoidance behavior. (A) Schematic of the imaging setup. (B) Individual (light blue) and fly-averaged (dark blue) calcium responses of LPLC1 population over time to a variety of visual stimuli (horizontally moving bars and squares, looming, square wave gratings, full-field flashes). Leftward in the stimulus schematics correspond to the back-to-front direction. (C) Time-averaged population responses of LPLC1 to horizontally translating bars and squares by the stimulus directions. Each dot represents an individual fly, and data from the same fly are connected by a gray line. (D) Cell-averaged spatial tuning curves of LPLC1 main dendritic stalks, measured with translating black squares. See also **Fig. S2B** for representative examples of calcium responses before averaging over time. (E) The vertical and horizontal receptive field sizes of individual LPLC1 dendritic stalks, measured as the full-width quarter-maximum visual angles of Gaussian fit to individual spatial tuning curves. (F) Time-averaged responses of individual LPLC1 cells to 10° x 10° black squares that passed through their receptive field centers. (G-J) Responses of individual LPLC1 cells to horizontally translating rectangular objects with various (G, H) heights and (I, J) widths, either as (G, I) functions of time by sizes or (H, J) peak responses as functions of sizes by directions. Time traces are only shown for the back-to-front directions. (K-M) LPLC1 cells responses to translating objects on rotating backgrounds, similar to behavioral results in **Fig. 1L-O**. (K) Responses over time to different object directions and background velocities. Vertical dotted lines and the shaded region respectively indicate the on-/offset of the background and the period during which the object was moving. (L) Peak calcium response as functions of background velocity, by the directions of the object. Positive velocity is in the same direction as front-to-back. (M) Differences of peak calcium responses between when the background was moving with and against the object, for each background speed. (N) Average direction selectivity index (DSI) of lobula dendritic ROIs of LPLC1 expressing jGCaMP7b, as a function of their estimated azimuthal receptive field center location. (O) The distribution of correlation between receptive field location and direction selectivity. Error bars and shading around mean traces all indicate standard error of the mean across flies (C, N, O) or cells (D-M). (B, C) N = 22 flies. (D-F) N = 80 (vertical), 60 (horizontal) cells. (G, H) N = 16 cells. (I, J) N = 12 cells. (K-M) N = 17 cells. (N, O) N = 37 flies. n. s.: not significant (p > .05); *: p < .05; **: p < .01; ***: p < .001; ****: p < .0001 in Wilcoxon signed-rank (C, M, O) or rank sum test (F).

To characterize the receptive field structure of LPLC1 neurons in more detail, we next recorded activity of individual LPLC1 neurons from their main dendritic stalks in lobula (**Fig. S2A**). For each cell, we first estimated their receptive field (RF) with translating black squares (Tanaka and Clark, 2020) (**Fig. S2B**; see *Methods*), and then subsequent stimuli were centered around the estimated RF location. On average, LPLC1 had a receptive field size of about 30° along both vertical and horizontal axes, measured as the full-width quarter-maximum value of the Gaussian fits (**Fig. 3E**). In addition, the response of LPLC1 neurons to stimuli used for RF mapping were significantly direction selective in the back-to-front and up directions (**Fig. 3F**).

We then measured the size tuning of LPLC1 by presenting horizontally translating rectangular objects with various heights and widths (**Fig. 3G-J**). This resulted in a tuning curve peaking at 10° of height (**Fig. 3H**), similar to several known lobula VPN types (Keleş and Frye, 2017; Städele et al., 2020; Tanaka and Clark, 2020). We confirmed that the LPLC1-dependent component of the slowing behavior is also tuned to small vertical sizes in an additional behavioral experiment (**Fig. S2C-F**). This was in contrast to slowing caused by LPC1 neurons, another back-to-front selective visual projection neuron (Eliason, 2017; Isaacson, 2018), revealing a complementary vertical size tuning between LPLC1 and LPC1 (**Fig. S2C-F**). On the other hand, LPLC1 was not tuned to objects with narrow width: rather, responses of LPLC1 increased up until the width of about 30° and saturated beyond that width (**Fig. 3I, J**). LPLC1 showed relatively broad tuning to object velocity and tuning for low flicker frequencies (**Fig. S2G-J**).

Next, we asked whether LPLC1 is itself sensitive to the relative motion between the objects and the background, as we found in the LPLC1-dependent slowing behavior (**Fig. 1L-O**). To test this, we measured LPLC1’s response to traveling squares over rotating checkerboard backgrounds similar to the stimuli used in the behavioral experiment (**Fig. 1M**). Overall, addition of moving background, especially fast ones, generally suppressed the response of LPLC1 neurons (**Fig. 3K, L**), similar to the behavioral slowing responses. Again, similar to the behavioral results, LPLC1 responded significantly more to back-to-front objects on backgrounds moving against rather than with the objects (**Fig. 3M**). This effect was weaker and not significant for front-to-back objects. This result suggests that the sensitivity to relative motion observed in the collision avoidance behavior is already computed at the level of LPLC1 calcium signals.

Lastly, we asked if the direction selectivity of LPLC1 population is spatially biased, a potential adaptation to the geometry of collisions (**Fig. 1C**) and a bias observed in the behavioral experiments (**Fig. 1P-R**). To this end, we recorded calcium responses in lobula dendrites of LPLC1 to rectangular objects sweeping long horizontal trajectories in either direction. We then calculated the direction selectivity of each dendritic ROI and plotted it against its estimated receptive field location (**Fig. 3N**). Direction selectivity of each ROI was quantified as direction selectivity index (DSI), calculated as the difference divided by the sum of its peak responses to stimuli moving in the front-to-back and back-to-front directions. Across flies, we found a strong correlation between the direction-selectivity of LPLC1 neurons and their azimuthal location (**Fig. 3N, O**), consistent with our earlier behavioral results. Thus, LPLC1 neurons are most direction selective in the regions of the visual field where direction is most predictive of a potential collision.

### LPLC1 receives inputs from T2, T3, and T4/T5

Having characterized physiological response properties of LPLC1 neurons, we next sought to obtain a mechanistic understanding how LPLC1 achieves these properties by combining its inputs. To identify neurons presynaptic to LPLC1, we turned to the hemibrain connectome dataset (Scheffer et al., 2020). First, we aimed to confirm the assumption that LPLC1 neurons receive inputs from T4/T5 tuned to back-to-front, upward, and downward motion at layers 2, 3, 4 of the lobula plate (i.e. T4/T5 subtypes b, c, and d) (Wu et al., 2016). While the hemibrain contains only a small fraction of lobula plate, it contains a large fraction of lobula as well as several labeled lobula plate tangential cells (LPTCs). Therefore, we hypothesized that we could still identify some T5 cells and examine their connectivity to LPLC1. Indeed, guided by their pre- and postsynapse innervations in lobula and lobula plate, connectivity to known LPTCs (or lack thereof), as well as their morphology, we were able to identify approximately 40 to 50 T5 cells in each of the four subtypes (**Fig. 4A, S3A**) (See *Methods* for details). See **Supplementary File 1** for the complete list of identified T5 cells. About 20% of the all identified T5b, c, and d cells synapsed onto identified LPLC1 cells, with the total synapse counts of about 50 per type (**Fig. 4B**). In contrast, we found only two synapses from T5a cells to LPLC1 (**Fig. 4B**). This observation supports the hypothesis that LPLC1 receives inputs from T5 at all layers of lobula plate it innervates (i. e., layers 2, 3, and 4). Beyond those anatomical connections, to confirm the functional connectivity between T4/T5 and LPLC1, we optogenetically activated the T4/T5 cells expressing Chrimson (Klapoetke et al., 2014) with a diode laser, while monitoring the axonal calcium activity of LPLC1 with jGCaMP7b. As expected, activation of T4/T5 resulted in large LPLC1 calcium transients in flies fed with ATR compared to negligible transients in control animals without ATR (**Fig. 4C, D**).

**Figure 4.**
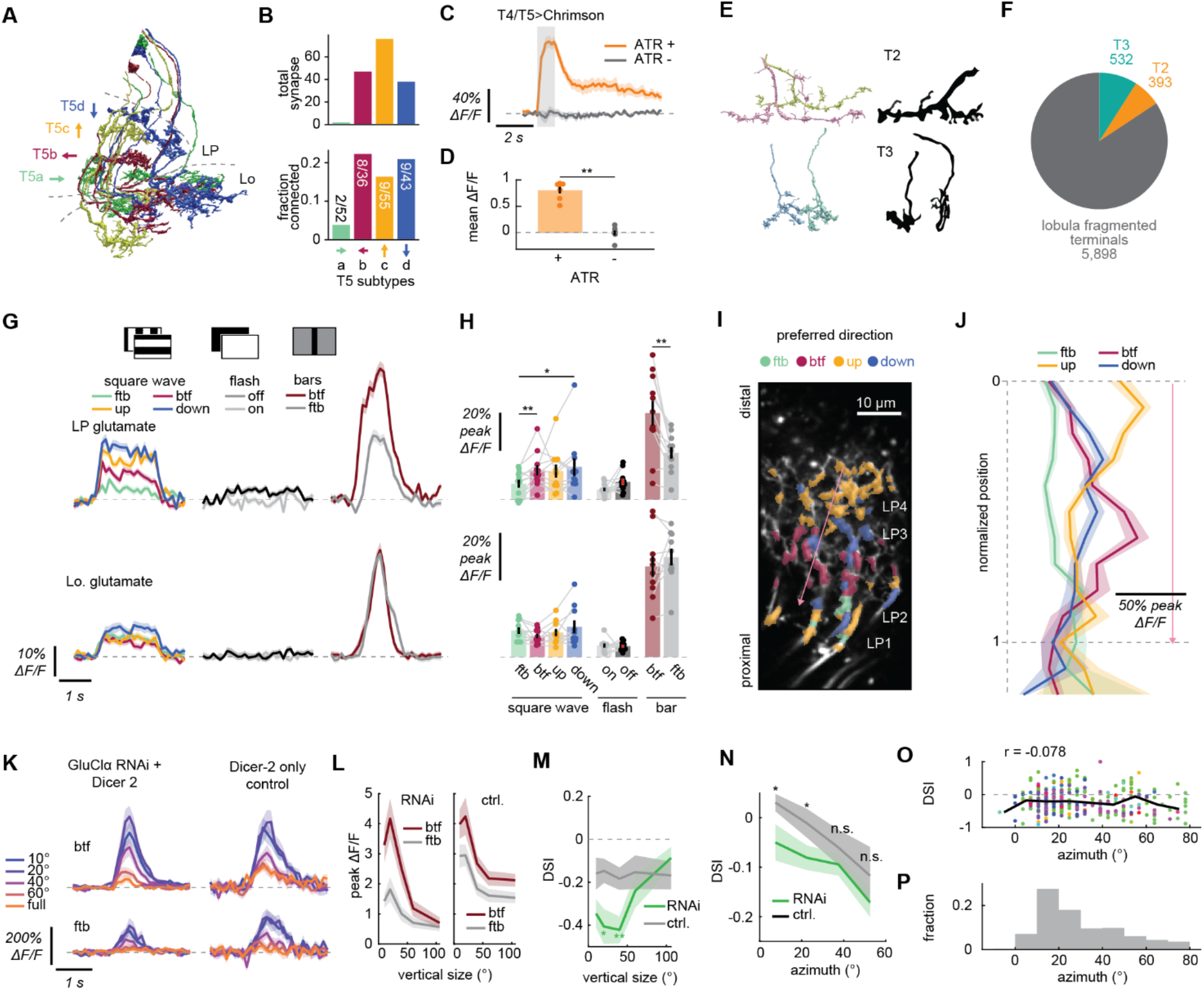
Input circuitry of LPLC1. (A) Examples of T5 cells in the hemibrain dataset, with the four subtypes coded by different colors. Characteristic layered innervation in lobul aplate (LP) and somata in lobula plate cortex are visible. See also **Fig. S3A**. (B) Connectivity from T5 cells onto LPLC1 by T5 subtypes, quantified by (*top*) the total number of synapses and (*bottom*) fraction of identified T5 cells connected to LPLC1. (C, D) Calcium response of LPLC1 to optogenetic stimulation of T4/T5 in flies with or without ATR feeding, either as (B) functions of time or (C) time-averaged. (E) Morphology of putative T2 and T3 axons from the hemibrain dataset (*left*), alongside with Golgi staining based morphology of T2 and T3 (*right*) (Fischbach and Dittrich, 1989). See **Supplementary File 3** for the list of visually annotated T2 and T3. (F) Total number of synapses the LPLC1 population in the hemibrain dataset receives from the putative T2 and T3 cells, among the other fragmented lobula terminals analyzed here. (G, H) Glutamate measured with iGluSnFR expressed in LPLC1 cells at (*top*) lobula plate (LP) and (*bottom*) lobula (Lo) dendrites to a variety of stimuli, either (E) over time or (F) time-averaged. (I) An example images of lobula plate dendrites expressing iGluSnFR, whose ROIs are color coded according to the direction of the bar to which they responded best. Approximate location of each lobula plate (LP) layer is indicated. The pink arrow indicates the axis along which we measured the normalized positions of ROIs in (J). (J) Peak glutamatergic signals in lobula plate dendrites, as functions of normalized positions of ROIs along the layers of lobula plate, measured from the distal most layer. (K-M) Calcium responses of LPLC1 cells expressing *GluClα* RNAi and their *Dicer-2* only controls to translating objects with various heights, as in **Figure 3G, H**. (K) Responses over time, by object sizes and directions. (L) Peak responses as the functions of object sizes, by object directions. (M) Direction selectivity index of the peak responses as the function of object size, by genotype. (N) Fly-averaged DSI of LPLC1 expressing *GluClα* RNAi with *Dicer-2* and their *Dicer-2* only control, as functions of azimuthal RF positions of the ROIs. (O) DSI of iGluSnFR signals in lobula plate dendrites in response to translating bars, plotted against the azimuthal RF location of each ROI. ROIs from different flies are in different colors, and the solid black line indicates median DSI within each 15° bin. DSI showed only weak correlation with the azimuthal location (r = -0.078). (P) The normalized histogram of azimuthal RF locations of lobula plate ROIs of LPLC1 expressing iGluSnFR. Error bars and shades around mean traces indicate standard error of the mean across flies, unless otherwise noted. (C, D) N = 7 (ATR+), 6 (ATR-) flies. (G, H) N = 11 (LP), 10 (Lo.) flies. (J, O, P) N = 17 (flies), 366 (ROIs). (K-M) N = 14 (*GluClα* RNAi), 16 (*Dicer-2* only) cells. (N) N = 21 (*GluClα* RNAi), 22 (*Dicer-2* only) flies. n. s.: not significant (p > .05); *: p < .05; **: p < .01; ***: p < .001; ****: p < .0001 in Wilcoxon signed-rank (H) or rank sum test (D, M, N). Non-significant pairs are not indicated in (H) for visual clarity.

Next, we tried to identify lobula neuron types providing excitatory inputs to LPLC1, specifically focusing on small-field columnar neurons. The hemibrain dataset does not contain most of the medulla neuropil. Thus the overwhelming majority of putative feedforward, columnar neurons that provide input to lobula (e.g., transmedullar (Tm) cells) are only partially reconstructed and are unlabeled. However, close inspection of their fragmented terminals can still offer useful insight into the input circuit organization of lobula VPNs (Tanaka and Clark, 2020). Here, we ran a connectivity- and morphology-based agglomerative hierarchical clustering on ∼1,000 fragmented terminals presynaptic to LPLC1, which likely represent feedforward excitatory inputs into LPLC1 and accounted for 25% of the lobula postsynapses in LPLC1 cells (**Fig S3B;** see *Methods* for details and **Supplementary File 2** for the complete results). Among the identified putative presynaptic cell types, of particular interest were T2 and T3 (Fischbach and Dittrich, 1989) (**Figs. 4E, S3B**). T2 and T3 are cholinergic (Konstantinides et al., 2018), small-field ON-OFF cells with tight size tuning, and they provide excitatory inputs to at least one other object-selective lobula VPN, LC11 (Keleş et al., 2020; Tanaka and Clark, 2020). We were able to identify 50 putative T2s and 82 putative T3s among the fragmented terminals analyzed here, which respectively had 393 and 532 total synapses on the entire LPLC1 population we analyzed of 60 cells (**Fig. 4F**). These numbers combined correspond to about one sixth of all synapses from the ∼1,000 small neurite fragments onto LPLC1 analyzed here (**Fig. 4F**). Overall, the connectomic analyses here suggest that LPLC1 achieves its direction selective response to small moving objects by pooling inputs from T2, T3, and T4/T5, among other neurons.

### Glutamatergic inhibition creates spatial bias in direction selectivity

Next, we wondered how the spatial bias in direction selectivity of LPLC1 could be implemented. One possibility is that inhibitory inputs are masking excitatory inputs from T4/T5 in a spatially biased manner. To characterize inhibitory inputs LPLC1 is receiving, we visualized glutamatergic signals at LPLC1 dendrites using iGluSnFR (Marvin et al., 2013). Glutamate is one of the major inhibitory neurotransmitters in the fly brain (Davis et al., 2020; Liu and Wilson, 2013), and several VPNs are known to receive directionally selective inhibition in lobula plate, including LPLC2 (Klapoetke et al., 2017; Mauss et al., 2015). We first presented flies expressing iGluSnFR in LPLC1 with a battery of visual stimuli consisting of full-field flashes, drifting square wave gratings, and vertical bars moving horizontally (**Fig. 4G, H**). We observed glutamatergic signals in both lobula and lobula plate neurites of LPLC1. Given that LPLC1 is cholinergic (Davis et al., 2020; Özel et al., 2020), these signals likely represent inputs into, rather than outputs from LPLC1. In both neuropils, the glutamatergic signals were strongest in response to the bars, moderate in response to the square waves, and minimal to the flashes (**Fig. 4G, H**). In addition, glutamatergic inputs in lobula plate, but not at lobula, were direction selective: in lobula plate, back-to-front bars elicited stronger glutamate signals than front-to-back ones. The front-to-back square wave also resulted in smaller responses than ones moving in the other three directions. Importantly, the direction selectivity of these measured glutamatergic signals is *syn-directional* with the preferred directions of LPLC1 itself and its excitatory inputs, unlike other VPNs that receive directionally opponent excitation and inhibition (Klapoetke et al., 2017; Mauss et al., 2015).

To better characterize this unexpected syn-directionally tuned glutamatergic inputs, we mapped the laminar organization of glutamatergic inputs into LPLC1 in the lobula plate. To do so, we presented the flies with vertical or horizontal bars translating in the four cardinal directions. Then, for each direction, we plotted the peak responses of dendritic ROIs against their relative position in the lobula plate along the distal-proximal axis (see *Methods* for details) (**Fig. 4I, J**). In the vertical directions, glutamatergic responses to upward motion peaked most distally (near layer 4), whereas responses to downward motion peaked slightly more proximally (near layer 3) (**Fig. 4J**). This observation is consistent with the previous documented innervation pattern and directional tuning of lobula intrinsic neurons LPi3-4 and LPi4-3, which are thought to receive excitatory input from one layer while providing glutamatergic inhibition in the neighboring layer (Mauss et al., 2015). In the horizontal directions, the peak of back-to-front responses was adjoining the peak of down responses proximally, likely corresponding to the layer 2 (**Figs. 4J**). The proximal-most ROIs (layer 1) showed more response to front-to-back bars than anywhere else (**Figs. 4J**), albeit with a smaller amplitude. This observation implies the existence of a glutamatergic interneuron types that receive inputs from T4/T5 in layers 1 or in layer 2 and send outputs locally within the same layer, in contrast to the LPi neurons studied previously (Mauss et al., 2015). We confirmed that this pattern of intra-layer glutamatergic inhibition in the horizontally selective lobula plate layers holds true beyond LPLC1 inputs by repeating the same experiment in flies expressing iGluSnFR pan-neuronally (**Fig. S3D, E**).

If these direction selective glutamatergic inputs into LPLC1 are indeed inhibitory, suppressing them should make LPLC1 more selective to back-to-front stimuli. To test this hypothesis, we knocked down a subunit of the glutamate-gated chloride channel *GluClα* specifically in LPLC1 by introducing RNAi (Liu and Wilson, 2013; Molina-Obando et al., 2019) while also overexpressing *Dicer-2*, which can facilitate mRNA cleavage (Kim et al., 2006). When we presented horizontally translating dark rectangular objects with various heights to the flies with RNAi, we observed that the responses of LPLC1 with *GluClα* RNAi to 20° and 40° tall objects were *more* selective for back-to-front direction compared to control genotype with only *Dicer-2* overexpression (**Fig. 4K-M**). This result confirms the idea that glutamatergic, syn-directional inhibition is suppressing the direction selectivity of wildtype LPLC1 neurons.

Finally, we tested whether this glutamatergic inhibition is responsible for the observed retinotopic bias in the direction selectivity of LPLC1. To this end, we again introduced *GluClα* RNAi and *Dicer-2* into LPLC1 and recorded population activity in lobula dendrites in response to objects moving horizontally. We found that the knock-down of *GluClα* significantly increased direction selectivity of forward-facing LPLC1 ROIs only (**Fig. 4N**). While the size of the effect was modest, this observation supports the idea that glutamatergic inhibition creates spatial bias of DSI in LPLC1. Conceivably, such bias can be inherited from glutamatergic neurons that already have spatially biased direction selectivity, or achieved *de novo* by the spatial bias in the synaptic strength between the glutamatergic neurons and LPLC1. To disambiguate these possibilities, we re-analyzed the iGluSnFR imaging data in lobula plate (**Fig. 4I, J**), and checked the distribution of azimuthal RF locations for ROIs and their direction selectivity. We found that the azimuthal location of ROIs did not correlate with their horizontal DSI (**Fig. 4O**), suggesting that the spatial bias in DSI is not simply inherited from the glutamatergic neurons. Interestingly, the majority of identified lobula plate ROIs in these iGluSnFR recordings had their RF centers in the frontal visual field (**Fig. 4P**). While this observation could simply reflect a bias in sampling, it could also favor the hypothesis that the spatial bias in the distribution of synapses between the glutamatergic neurons and LPLC1 is creating the bias in direction selectivity.

### A downstream pathway that mediates collision avoidance faithfully inherits LPLC1’s response

In a last set of experiments, we aimed to identify pathways downstream of LPLC1 that transmit signals responsible for the collision avoidance behavior. We focused our experiments on five major neuron types postsynaptic to LPLC1: DNp03, DNp06, PVLP112/113, and PLP219 (**Figs. 5A, S4A**), which could be selectively labeled by split Gal4 lines (Namiki et al., 2018), including ones we newly generated (see *Methods* for details) (**Figs. 5B, S4B**). These five cell types accounted for more than half of total LPLC1 outputs (approximately 9,500 out of 17,000 total synapses) and about 70% of total central brain outputs of LPLC1 (approximately 14,000 synapses) (**Fig. S4C**). Of these neurons, the two descending neuron types, DNp03 and 06 were promiscuous in receiving inputs from VPNs. In addition to LPLC1, DNp03 receive inputs from LPLC4 and LC4, and DNp06 from LC4, 6, and 31. In contrast, the interneurons PVLP112/113 and PLP219 receive about one half of their inputs from LPLC1. We treated PVLP112 and 113 as a single group, because they share very similar connectivity and morphology, and our split Gal4 line appeared to label both, based on the number of cell bodies (4 and 3 PVLP112 and 113 are respectively reported in the hemibrain dataset, and the split Gal4 line typically labeled 7 PVLP cells per hemisphere) (**Fig. S4A, B**).

**Figure 5.**
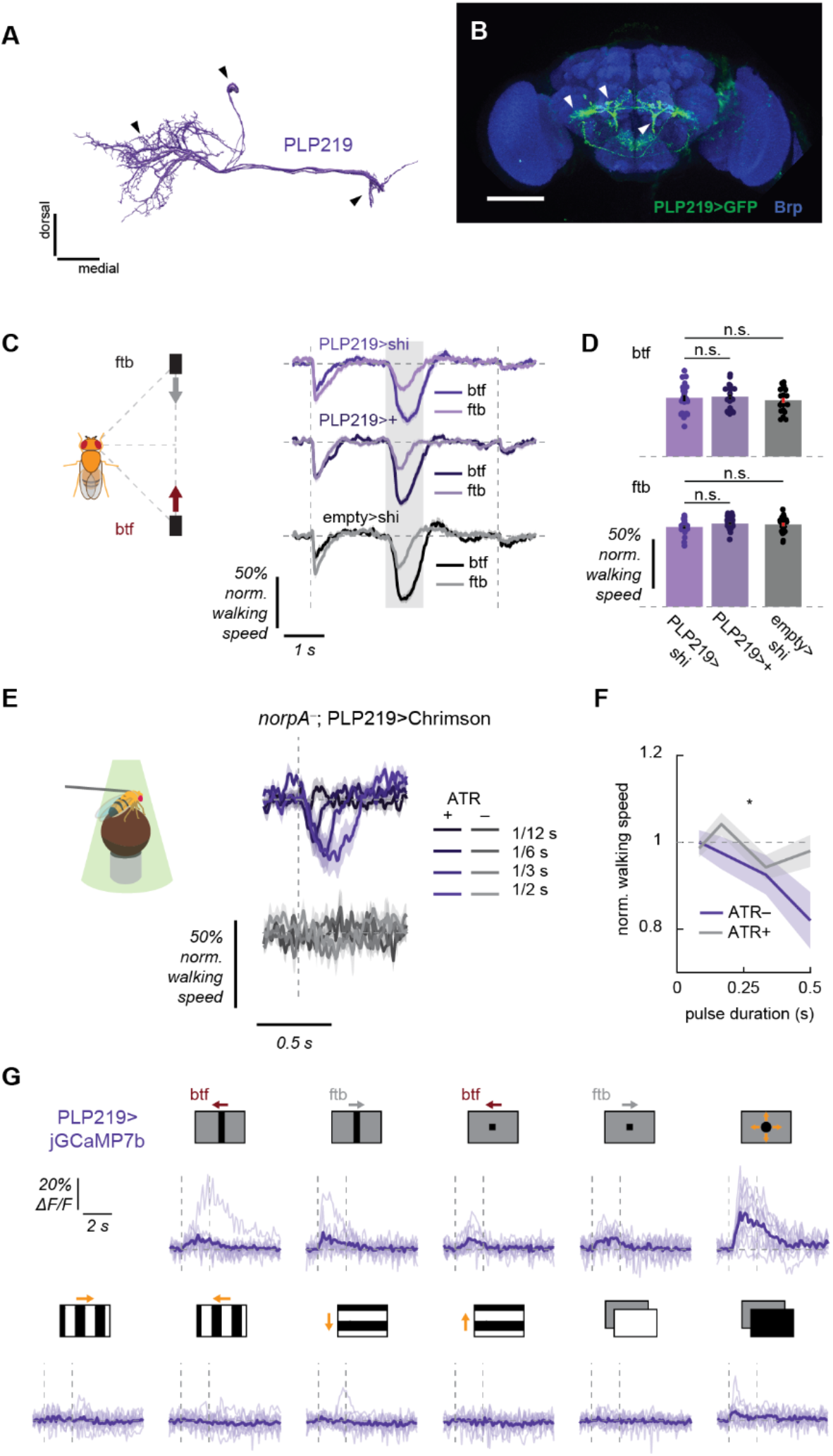
A central brain pathway for collision avoidance. (A) Reconstructed morphology of PLP219 neurons, viewed from the front. (B) The expression pattern of a newly generated PLP219 split Gal4 line (VT041832AD, VT021792DBD > UAS-myr::GFP). The corresponding structures are marked by the arrows between (A) and (B). (C, D) Walking responses of PLP219 silenced flies and their controls in response to the parallel stimuli, (C) over time or (D) averaged over time. (E, F) Walking responses of PLP219>Chrimson flies with or without ATR feeding in response to pulses of green light with different durations, either (C) over time or (D) time-averaged. (G) Individual fly (light purple) and averaged (dark purple) calcium responses of PLP219 population over time to a variety of visual stimuli (horizontally moving bars and squares, looming, square wave gratings, full-field flashes). Leftward in the stimulus schematics correspond to the back-to-front direction. Error bars and shades around mean traces all indicate standard error of the mean. (C, D) N = 20 (PLP219>shi), 19 (PLP219/+), 22 (empty/shi). (E, F) N = 15 (ATR+), 13 (ATR-). (G) N = 11. n. s.: not significant *: p < .05 in (D) Wilcoxon signed-rank test or (F) 2-way analysis of variance (ANOVA) (the main effect of ATR conditions).

To test whether any on these downstream neurons is necessary or sufficient for the collision avoidance behavior, we repeated synaptic silencing and optogenetic activation experiments identical to those we performed for LPLC1 (**Fig. 2**). Somewhat surprisingly, given how these four neuron types receive the majority of LPLC1 outputs, silencing of none of the four with *shibire^ts^* resulted in any significant change in slowing response to the parallel stimuli in either direction (**Figs. 5C, D, S4D, E**). In contrast, optogenetic activation of PLP219 with Chrimson caused flies to slow significantly (**Figs. 5E, F, S4F, G**), similar to activation of LPLC1 (**Fig. 2G-I**). The results show that activity of PLP219 is sufficient to trigger slowing in the absence of visual inputs, while its output is not necessary and is likely redundant with other parallel pathways, which could also include neurons that we did not include in the present survey.

Finally, to characterize the visual response properties of PLP219 neurons, we imaged the calcium activity of their putative dendrites with jGCaMP7b (**Fig. 5G**) while presenting the same broad battery of stimuli we used for the initial glomerular imaging of LPLC1 (**Fig. 3B**). Overall, the pattern of PLP219’s responses closely matched those of LPLC1 axon terminals, where they responded to moving bars, squares, and expanding discs, but not to full-field stimuli (**Figs. 3B, 5G**). This was in contrast to PVLP112/113 neurons, which responded to a broader set of stimuli, including drifting gratings and flashes (**Fig. S4H**). In summary, PLP219, a downstream pathway of LPLC1 that mediates collision avoidance, inherits the response property of LPLC1 faithfully than another parallel pathway.

## Discussion

In the present study, we explored a collision avoidance behavior in walking *Drosophila* and its underlying circuit mechanisms as a simple model of motion-based spatial vision. Using high-throughput psychophysics experiments, we demonstrated that back-to-front motion in the frontolateral visual field—a geometrical cue for near collision—causes slowing in walking flies (**Fig. 1**). Using genetic silencing and activation experiments, we showed that the visual projection neuron LPLC1 is necessary for this putative collision avoidance behavior and its activity is sufficient to cause slowing in walking flies (**Fig. 2**). Physiological response properties of LPLC1 mirrored the visual tuning of the slowing behavior, most notably in its spatial bias in direction selectivity (**Fig. 3**), which was also consistent with the geometry of near collisions. Using connectomic analyses, optogenetics, and neurochemical imaging and manipulation, we showed that object-selective T2 and T3 inputs are pooled with direction-selective T4/T5 inputs, likely establishing the object- and direction-selectivity of LPLC1, while spatially biased glutamatergic inhibition creates its position-dependent tuning (**Fig. 4**). Lastly, we identified a downstream neuron of LPLC1 called PLP219 to be sufficient to cause slowing, and to inherit the response property of LPLC1 faithfully (**Fig. 5**).

### Positional cues for threat detection and collision avoidance

As objects move relative to an observer, the apparent size and position of the object systematically change as dictated by geometry. How animals detect change in object size and use it to avoid predation has been well studied in various vertebrate species ranging from primates (Schiff et al., 1962), rodents (De Franceschi et al., 2016; Kim et al., 2020; Yilmaz and Meister, 2013), birds (Sun and Frost, 1998), and fish (Temizer et al., 2015), as well as in insects (Card and Dickinson, 2008; Gabbiani et al., 1999, 2002; Klapoetke et al., 2017; von Reyn et al., 2014). In contrast, less is known about how and when animals use positional changes or directional motion to detect and avoid collision with moving objects. In general, positional changes of moving objects are more salient than their changes in apparent size: One can show that the maximum apparent expansion rate of an object with radius *R* moving at a given speed is always less than its maximum apparent translational velocity when the object is more than *R* away from the observer (see *Methods* for calculation). Moreover, the ratio between the maximal translation rate and the maximal expansion rate can become arbitrarily large as the object is further and further from the observer (see *Methods* for calculation). Intuitively, these results correspond to the fact that one can easily tell whether someone 100 meters away is running to the right or left, while it is difficult to tell if that same person is running towards or away from you, based solely on visual motion. This saliency of translation rates is likely one reason that aerial predators employ interception strategies that minimize their apparent positional shifts on their prey’s retinae (Ghose et al., 2006; Kane and Zamani, 2014; Mischiati et al., 2015). Less sophisticated pursuit strategies, often used in non-predatory chasing among conspecifics (Chiu et al., 2010; Land, 1993), generate positional changes that can be used by pursuees to detect pursuers. Note that even predators that employ sophisticated strategies will suffer from positional changes after sudden turns of the prey until they settle into a new interception course.

Positional changes are therefore a useful cue to simply detect objects such as conspecifics and predators, but back-to-front motion in particular can be predictive of future collisions. This is because approaching objects appear to be moving back-to-front only when they will cross the path of the observer in front, which would then pose collision risks if the object slows or stops. We empirically confirmed this conjecture by running a simple simulation with randomized trajectories (**Fig. 1C**). Based on this geometrical argument, we interpret the direction selective slowing behavior of the flies studied here as a collision avoidance behavior. This is in contrast to other object motion-triggered freezing behaviors in both flies (Tanaka and Clark, 2020) and mice (De Franceschi et al., 2016), which are not selective for stimulus direction and thus are unlikely to be a specific response to predicted collision.

### Retinotopic bias in LPLC1 matches the geometry of collisions

In this study, we found retinotopic biases in the direction selectivity of both behaviors and neural processing. First, the direction selectivity of the collision avoidance slowing to the back-to-front direction was more pronounced in the frontolateral visual field. In addition, direction selectivity of LPLC1 neurons also strongly correlated with the azimuthal location of their receptive field. Since the frontolateral visual field is where back-to-front motion is most predictive of immediate collision (**Fig. 1C**), the spatial bias in the LPLC1 circuitry can be seen as an adaptation to this geometry.

Retinotopic biases in visual processing have been found in diverse species. For example, in vertebrate retinae, circuit features such as opsin expression, dendritic morphology, and synaptic strengths can all vary systematically across visual space, depending on species (Bleckert et al., 2014; Heukamp et al., 2020). It is also well established that features such as receptive field sizes (Harvey and Dumoulin, 2011) and orientation selectivity (Sasaki et al., 2006) exhibit retinotopic biases in primate visual cortices. Although these biases have been variously speculated to be adaptations to unique sensory ecology of different species, few were connected to strong geometrical explanations or to behavior. Importantly, the geometrical justification we provided here for the spatial bias in direction selectivity for collision detection is not specific to flies. Thus, it is likely that similar biases exist in other sighted species, arrived at through convergent evolution. Indeed, rodent superior colliculus—a center of visual threat detection—has been reported to exhibit a similar retinotopic bias where back-to-front and upward motion is overrepresented in the upper lateral visual field (Li et al., 2020), likely mirroring the geometry of approaching overhead predators.

### Other behavioral functions of LPLC1 neurons

Although here we focused on LPLC1’s involvement in collision avoidant slowing behavior in walking flies, this does not preclude the possibility that LPLC1 is involved in different behavioral programs in other sensory and behavioral contexts. Supporting this idea, we found multiple downstream neurons of LPLC1 whose activation did not result in slowing and also had divergent visual response properties. Indeed, a previous study reported that strong optogenetic activation of LPLC1 can lead to behavioral phenotypes other than slowing, such as jumping (Wu et al., 2016). Descending neurons DNp03 and DNp06, which receive inputs from other loom-sensitive, jump-inducing VPNs (LC4, LC6), make good candidates for the neural basis of such jumping phenotypes.

An interesting question is how the activation of LPLC1 neurons by different stimuli (e. g., small objects moving back-to-front vs. looming objects) results in different behavioral responses. For example, one can imagine that the activation of LPLC1 without activation of other loom sensitive cells (e.g., LC4, LC6) is decoded as the presence of a conspecific in a collision course to initiate slowing, whereas simultaneous activation of LPLC1 alongside other loom detectors strongly implies predators and thus triggers rapid escape. How such population-level decoding and behavioral decision-making is implemented through the network of interglomerular local neurons (Mu et al., 2012) is of particular interest for future studies.

### Convergence of motion and object detectors

In flies, the lobula complex consists of the lobula and lobula plate, which are the highest order brain neuropils that remain specialized for visual processing. Among these two neuropils, lobula plate has been historically under intensive study as the neural basis of visual motion detection and stabilization reflexes (Hausen, 1976; Maisak et al., 2013), while the functions of the lobula neuropil have remained less clear. The recent series of studies on lobula output neurons (Keleş and Frye, 2017; Klapoetke et al., 2017; Morimoto et al., 2020; von Reyn et al., 2017; Ribeiro et al., 2018; Städele et al., 2020; Tanaka and Clark, 2020; Wu et al., 2016) have started to show that these neurons detect ethologically relevant objects, like mates and predators, to drive specific behavioral programs. Visual projection neurons innervating both lobula and lobula plate, including LPLC1, are uniquely situated to integrate these object and motion signals. Here, we showed that LPLC1 likely pools inputs from motion- and object-detecting interneurons (T4/T5 and T2/T3 neurons) to construct a more complex visual feature. While there are other visual projection neuron types spanning lobula plate and lobula whose physiology have been studied (for instance, LPLC2 (Klapoetke et al., 2017), LLPC1 (Isaacson, 2018)), lobula inputs to those neurons remain to be explored.

Interestingly, a similar computational motif of convergence between motion- and object-detecting pathways seems to be present in the early visual systems of vertebrates as well. Vertebrate retinae are equipped with retinal ganglion cells selective for motion directions (Barlow and Hill, 1963) as well as small objects (Ölveczky et al., 2003; Semmelhack et al., 2014; Zhang et al., 2012). The axon terminals of motion- and object-selective ganglion cells innervate shallowest layers of optic tectum in zebrafish (Robles et al., 2014) as well as of superior colliculus in mice (Hong et al., 2011). While the internal circuitry of the optic tectum / superior colliculus is still not well understood, physiological studies on the neural bases of prey capture in larval zebrafish have identified tectal neurons that show direction selective responses to small objects similar to LPLC1 (Antinucci et al., 2019; Bianco and Engert, 2015; Förster et al., 2020). Similarly, narrow field neurons in mouse superior colliculus, which are also necessary for prey capture behavior, exhibit direction selectivity as well as tight tuning to small object sizes (Hoy et al., 2019). These results suggest that integration of motion- and object-detector outputs similar to LPLC1 indeed takes place in the optic tectum / superior colliculus. Parallels between vertebrates and invertebrates in the early layers of visual processing and motion detection have been noted (Borst and Helmstaedter, 2015; Clark and Demb, 2016; Sanes and Zipursky, 2010). The findings reported here extend the computational analogies between insect and vertebrate visual systems to the motif of initial segregation and subsequent convergence of motion and object detecting pathways to drive specialized object-detection behaviors.

## Acknowledgements

We thank the members of the Clark lab as well as A. Nandy, J. Jeanne, and L. Liang for helpful comments and discussions. RT was supported by the Takenaka Foundation and the Gruber Foundation. DAC and this project were supported by NIH R01EY026555, NIH R01NS121773, and NIH P30EY026878.

## Methods

### Resource Table

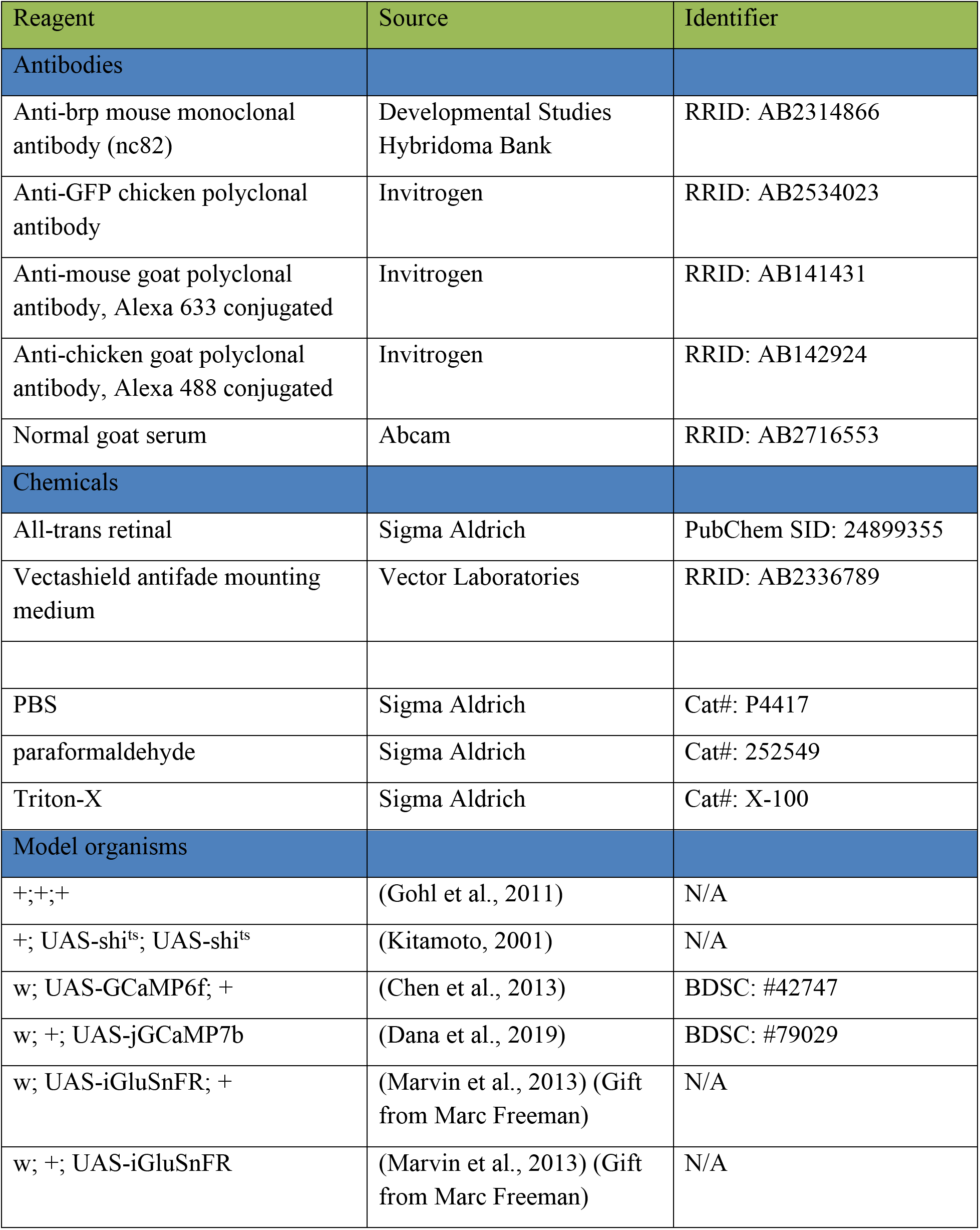
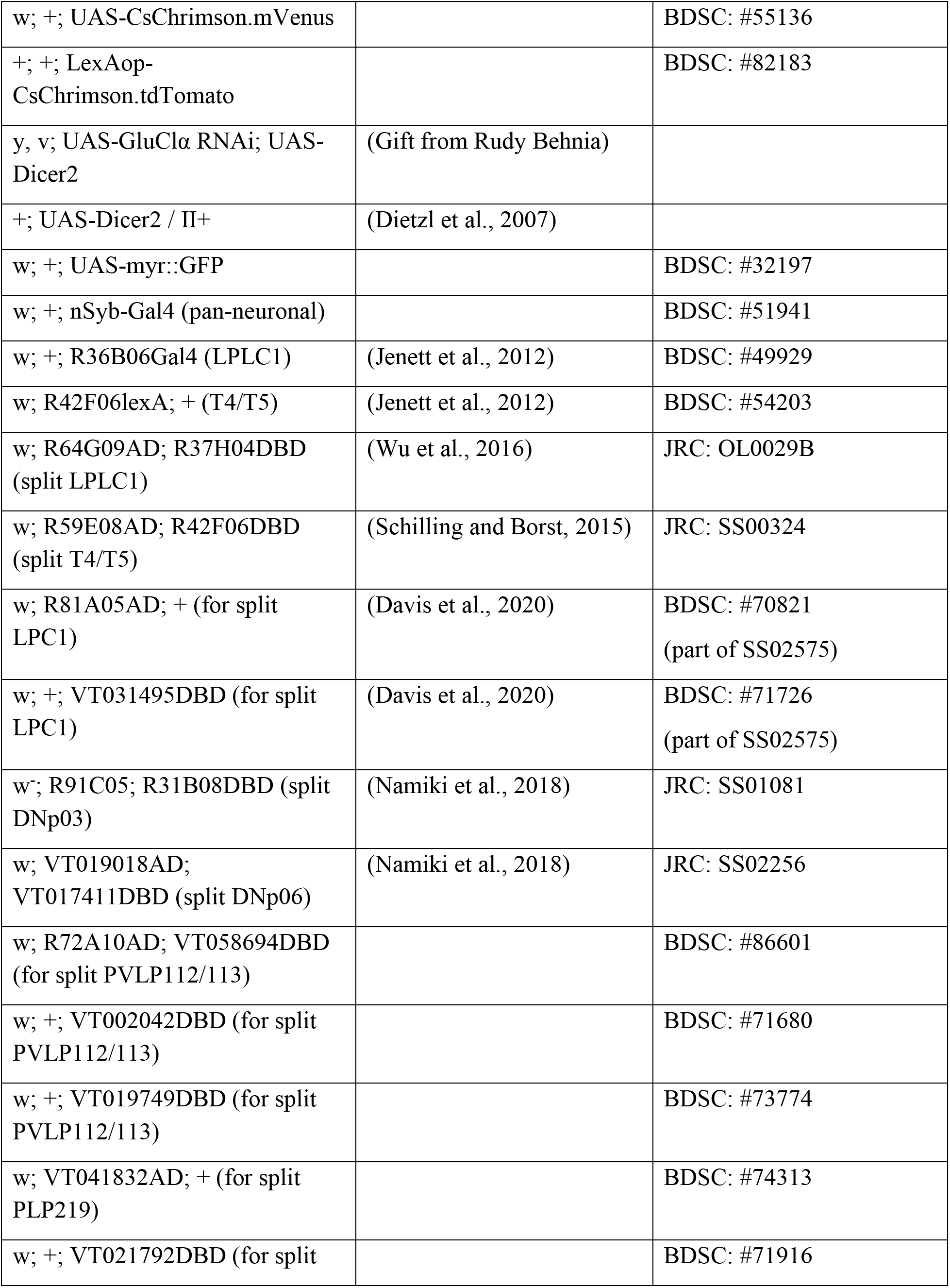
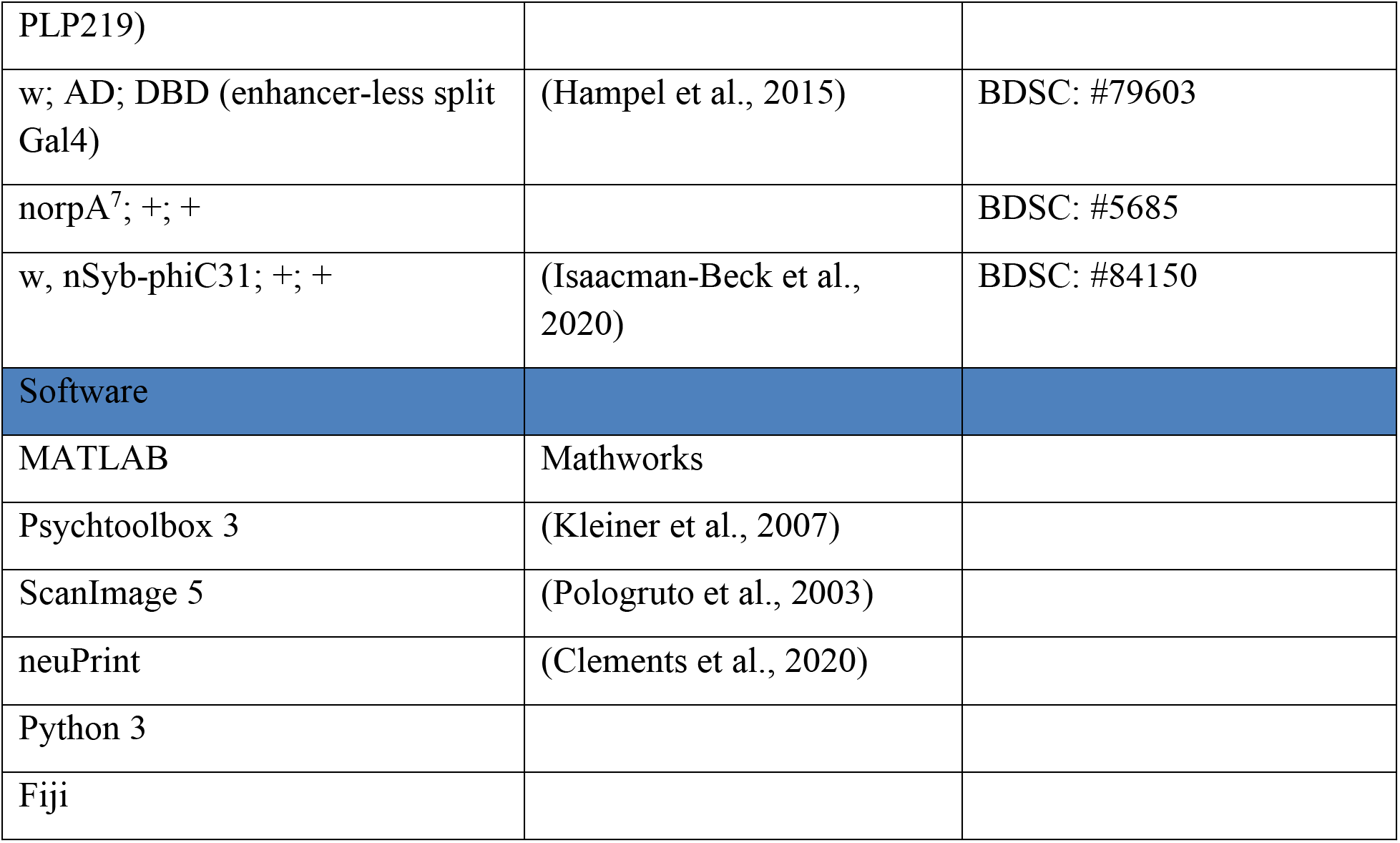

### Fly strains and husbandry

Flies were raised at around 50% humidity on a dextrose-based food. Non-virgin female flies were used for all experiments except for the optogenetic activation in blind flies, where male flies with single deficient allele of *norpA* on the X chromosome were used for experimental convenience. Flies for behavior experiments were raised at 20 ℃ on 12 h light/dark cycle. Adults less than one day post eclosion were collected with CO_2_ anesthesia, and all experiments were performed within 12 to 24 h after staging, with the exception of flies for optogenetics experiments, which were dark-reared on food with or without 10 µM all-trans retinal (ATR) (de Vries and Clandinin, 2013). All behavioral experiments were performed within 3 h windows after lights-on or before lights-off. Flies for imaging experiments were grown at 25 ℃. Most flies were staged with CO_2_ at least 12 h prior to the experiments and immobilized with ice before surgery. Flies were typically imaged between 2 to 7 days post eclosion. Flies for imaging experiments with optogenetics were dark reared on food with or without ATR for 3 days. In imaging experiments with RNA interference, only 5 days old flies were used. The genotypes of the flies used for the experiments are summarized in **Supplementary Table 1**.

### Tethered walking psychophysics assay

Previously reported fly-on-the-ball rigs were used to measure fly locomotor responses to visual stimuli (Creamer et al., 2019). Flies were anesthetized on ice, and tethered to 30G surgical needles with UV-curable epoxy on their dorsal thorax. The tethered flies were mounted above air-floated balls, whose rotation were used as a read out of flies’ attempted movements. The rotation of the balls was measured with optical mouse chips at the resolution of ∼0.5° and 60 Hz. Visual stimuli were projected onto panoramic screen covering 270° azimuth and 106° elevation using Lightcrafter DLP evaluation module (Young Optics) using green light (peak 520 nm and mean intensity ∼ 100 cd/m^2^). The temperature of the rig was set at 36 ℃ to promote walking and to use thermogenetic tools.

Visual stimuli used in the behavioral experiments were compiled in **Supplementary Table 2**. For optogenetic stimulation (Creamer et al., 2019), the panoramic screens were removed and the pulses of green light were directly shone on the flies from the four directions (top, front, left, right). The mean light intensity was approximately ∼10 µW/mm^2^.

### Behavioral data analysis

Walking speed of the flies were normalized relative to the average walking speed within the 500 ms window prior to each stimulus onset, unless otherwise noted. The time traces of normalized walking speed and turning angular velocity were then averaged across presentations of each stimulus type. Walking and turning time traces in response to mirror-symmetric pairs of stimuli were also averaged in subtractive and additive fashion, respectively. These individual mean time traces were then averaged over time for statistical comparisons. The window for the averaging spanned the entire duration of stimuli, unless otherwise noted in the caption. In addition, group mean time traces and standard error of the mean were calculated from the individual mean time traces to visualize the dynamics of the responses.

### Two-photon imaging

For imaging experiments, flies were cold anesthetized and head-fixed into a metal shim with UV curable epoxy. The brain was exposed by surgically removing cuticle, fat tissue, and trachea on the back of the head. All recordings were performed on the right side of the brain. The mouth parts were fixed with the epoxy to minimize the brain movement. The exposed brain was submerged under oxygenated sugar-saline solution (Wilson et al., 2004). Imaging was performed with a two-photon microscope (HyperScope; Scientifica) equipped with a 20x water immersion objective (XLUMPlanFL; Olympus). Visual stimuli were presented on a panoramic screen covering 270° azimuth and 69° elevation of the flies’ visual field with a DLP projector (Texas Instruments) (Creamer et al., 2019). Stimuli were pitched 45° forward relative to the screen to account for the tilt of the fly’s head in the shim. The projector output was filtered with a 565/24 in series with a 560/25 filter (Semrock) to prevent green light from bleeding into the PMT. The input into PMT was also filtered with two 512/25 filters (Semrock) to capture green fluorophore emissions. A femtosecond Ti-sapphire laser (Mai Tai; Spectra-Physics) provided 930 nm excitation. The power on the sample was set below 40 mW. Images were acquired at 8.46 Hz with ScanImage (Pologruto et al., 2003) software and motion-corrected offline. Frames with more than 4.3 microns of motion were excluded from further analyses, and recordings with more than 5% of frames rejected were discarded.

### Stimulus presentation

The stimuli used in behavioral and imaging experiments are respectively compiled in **Supplementary Tables 2** and **3**. In some imaging experiments, probe stimuli (**Supplementary Tables 4**) were presented at the beginning of experiments in order to identify responsive ROIs. See the section on imaging data analysis for how responses to probe stimuli were used in the analysis. All visual stimuli were presented against mean gray background unless otherwise noted. Visual objects were all black and presented on the visual equator unless otherwise noted. Each stimulus presentation was interleaved with blank gray screen, typically around 3 s. When a stimulus is described in terms of azimuthal and elevational degrees, the azimuthal and elevational zero respectively correspond to the central meridian and visual equator, with positive degrees indicating right and ventral visual fields. Since all the imaging experiments were performed on the right hemisphere, positive horizontal velocity always corresponds to front-to-back movements. Stimuli used in the single cell imaging experiments (**Figs. 3G-M, 4K-M, S2G-J**) are centered about the estimated receptive field location of the recorded cell.

### Optogenetic activation during imaging

Optogenetic activation of Chrimson under the two-photon microscope (**Fig. 3C, D**) was performed using a Thorlabs 690 nm laser diode (Thorlabs, HL6738MG). The measured power of the laser at the sample was ∼2 mW/mm^2^ and the laser was shone onto the sample through the imaging objective.

### Imaging data analysis

ROIs were defined either manually (glomerular and single-cell recordings; **Figs. 3B-M, 4C, D, K-M, S2AB**), with a watershed segmentation algorithm (Meyer, 1994) (dendritic recordings; **Figs. 3N, O**, **4G-J, N-P, S3C, 5G, S4H**) based on time-averaged fluorescent images, or as 3 µm rectangular grids (pan-neuronal recordings; **Fig. S3D**). To remove stimulus bleed-through, the recordings were subtracted with the pixel-averaged signals from background regions, which were defined as the largest contiguous regions below 10 percentile brightness. The fluorescent time traces were then converted into the unit of ΔF/F to account for expression level variability and photo-bleaching of the fluorophores. To obtain the baseline fluorescence (*i.e.*, the denominator F), fluorescence within each ROI was averaged across pixels, and a decaying exponential *Ae^-τ^* was fit to the time-averaged fluorescence within each interleave epoch, where τ was constrained to be identical across all ROIs in a single recording. The fit exponential (*i.e.*, the baseline fluorescence) was then subtracted from the original ROI-wise fluorescence time traces, and the remainder (ΔF) was then divided by the same fit exponential to generate ΔF/F time traces.

In some recordings where ROIs were extracted in an automated fashion (**Figs. 3N, O**, **4G-J, N-P, S3D, E**), responsive ROIs were selected based on the consistency of their responses to probe stimuli (see **Supplementary Table 4**). The probe stimuli were typically presented three to five times before each recording, and Pearson correlations between every pair of responses were calculated. ROIs with average correlation below certain thresholds were then discarded (0.4 for GCaMP6f and jGCaMP7b recordings, 0.3 for iGluSnFR recordings).

The responses to repetitions of the same stimulus were averaged within each ROI, and then across all ROIs within each fly to generate an individual mean response. The time-averaged ΔF/F during the 500 ms period preceding each stimulus presentation was subtracted from the time trace to remove the spontaneous fluctuation of ΔF/F. For statistical comparisons across conditions and genotypes, mean or peak individual mean responses were calculated over appropriate time windows, which spanned the entire duration of the stimuli unless otherwise noted. Additionally, group mean responses and standard error of the mean were calculated based on the individual mean responses across flies to visualize the dynamics of the responses.

In some lobula plate recordings (**Figs. 4I, J, S3D, E**), the laminar positions of ROIs were estimated. To this end, we manually drew a directed line segment that approximately started at the distal end of lobula plate, traversed the layers orthogonally, and ended at the proximal end. The position of each ROI along this line segment was calculated as a proxy of its layer affiliation.

### Receptive field localization

In the single cell recordings (**Figs. 3G-M, S2G-J**), the receptive field (RF) location of each cell was mapped prior to the experiment, and subsequent stimuli were centered around the estimated RF location (Tanaka and Clark, 2020). First, the approximate RF location was probed interactively by presenting translating small black squares. Next, a 10° black square moving horizontally or vertically at 60 °/s swept the 40° x 40° area around the approximate RF location at the resolution of 5° (noted as RF mapping stimulus in **Supplementary Table 3**). For each azimuth and elevation, the neural response in the unit of ΔF/F (see later) was averaged over time within the 1.5 s window from the stimulus onset and over the directions of motion, resulting in horizontal and vertical spatial tuning curves. Gaussian functions were independently fit to the two tuning curves, and resulted means of the distributions were used as the estimated RF center. In addition, the full-width quarter-width (FWQM) values of the fitted Gaussian functions were later used as the measure of RF size (**Fig. 3E**). Only sizes of RF with good (R^2^ > 0.8) Gaussian fit are plotted for this purpose. In some non-single cell dendritic recordings (**Figs. 3N, O**, **4N-P**), azimuthal RF location of each ROI was estimated based on the averaged time-to-peak in response to objects moving rightward and leftward.

### Geometrical simulation

For the simulation in **Figure 1C**, 5 million circular objects with 2 mm radius were simulated around an observer. The positions of the objects were uniformly distributed within a circular area with the radius of 200 mm about the observer. We assumed the observer to be moving forward at 10 mm/s, and the speed of the objects were randomly drawn from a uniform distribution ranging from 0 to 20 mm/s. The direction of the objects’ velocity was also chosen uniformly at random. For each object, given the instantaneous relative position and velocity and under the assumption that the both observer and the object maintain the constant velocity, we calculated immediate collision risk as time-discounted, rectified inverse intercept between the observer and the object trajectories. The intercept *I* and immediate collision risk *h* are given as follows:

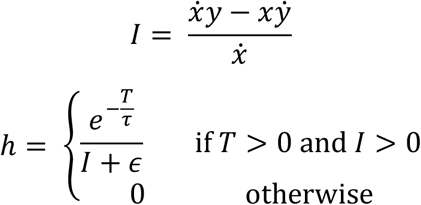

where (*x*, *y*) and (*ẋ*, *ẏ*) are the initial position and velocity of the object relative to the observer, *T* = −*x*/*ẋ* (time to path crossing), *τ* = 10 s and ϵ = 2 mm. We then plotted *h* as a function of the instantaneous angular position and velocity of the object as seen by the observer, averaged over samples. The code to run the simulation is available on GitHub (https://github.com/ClarkLabCode/CollisionSimulation)

### Proof of the geometrical conjecture

Let us assume a stationary observer at the origin and a circular object with radius *R* located at **r_0_** = [x_0_, y_0_], moving at a constant velocity **v** = [v_x_, v_y_]. Let us denote the future position of the object as **r**(t) = [x_0_ + v_x_t, y_0_ + v_y_t] = **r_0_** + **v**t, and distance to the object d(t) = |**r**(t)|. Then, the future retinal position φ(t) and size ψ(t) of the object seen from the observer can be written as

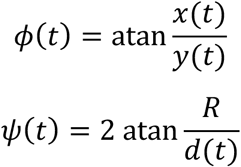

where φ = 0 points in the positive direction along y-axis. Then, the instantaneous angular velocity and expansion rate at time t = 0 can be obtained by differentiating these by t and evaluating at t = 0:

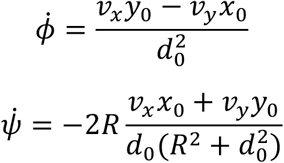

where d_0_ = d(0). Now, if we constrain the speed of the object to be a constant v = |**v**|, v_x_ and v_y_ can be written as [v_x_, v_y_] = v[cosθ, sinθ], where θ is the direction of the object’s movement. We can also set x_0_ = 0 and y_0_ = d_0_ without losing generality Then, maximum angular velocity and expansion rate of the object are

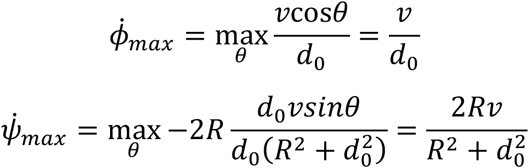

The ratio between these two values can be written as

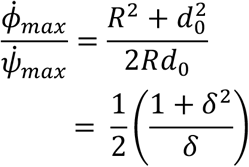

where *δ* = *R*/*d*_0_. This is a monotonically increasing function of δ that grows arbitrarily large with *δ*, which is larger than 1 when *δ* > 1. That is, when the object is further than *R* away from the observer, its apparent angular velocity caused when it moves tangentially to the observer is larger than the expansion rate caused when it moves straight toward the observer.

### Connectomic identification of T5

To identify candidate T5 cells in the hemibrain v1.1 dataset (Scheffer et al., 2020), we first extracted cells that had (1) synapses only in lobula or lobula plate, and (2) more presynapses in lobula plate than in lobula and more postsynapses in lobula than in lobula plate. From this candidate T5 pool, we identified cells that were connected to only one out of the three sets of monostratified LPTCs in single lobula plate layers (HS and CH for layer 1, H2 for layer 2, and VS for layer 4, respectively). After visual inspection, we were able to identify 52 T5a, 36 T5b, and 43 T5d cells. Since there is no identified monostratified LPTC in layer 3 of lobula plate, we searched for T5c cells from the candidate T5 pool as ones that had (1) no connection to the aforementioned LPTCs, and (2) fewer pre- and postsynapses in lobula and lobula plate than the corresponding maximum numbers of pre- and postsynapses in the two neuropils among the T5a, b, d cells identified above. After visual inspection, this resulted in 55 T5c cells (**Figs.4A, B, S3A**). We then examined the connectivity between the identified T5 subtypes and LPLC1. The code to identify candidate T5s can be found on our GitHub repository (https://github.com/ClarkLabCode/LPLC1ConnectomeAnalysis), and the body IDs of the annotated T5s can be found in **Supplementary File 1**.

### Connectomic identification of lobular inputs into LPLC1

To identify columnar neuron types providing inputs into LPLC1 neurons in the lobula, we first extracted all neurons in the hemibrain v1.1 dataset (Scheffer et al., 2020) that have (1) at least 3 synapses onto a single LPLC1 neuron, (2) no synapse outside of lobula, and (3) less than 300 synapses in total, pre- and postsynapses combined. This resulted in a pool of 977 distinct lobula intrinsic terminals. We then clustered these lobula intrinsic terminals according to their (1) connectivity, (2) terminal morphology, and (3) layer innervation patterns. First, we identified all labeled cell types that had more than 2 synapses from at least a single cell among the pool of the lobula intrinsic terminals, which resulted in 126 distinct identified cell types, including LPLC1. We then constructed a 977 x 126 matrix that contained synaptic counts between each lobula intrinsic terminal and each postsynaptic cell type. Postsynaptic cells without identified cell types were ignored here. Second, we extracted the positions of the all presynapses of each lobula intrinsic terminal in the native XYZ coordinate of the hemibrain dataset. Then, these synapse positions were translated and rotated such that the new XY axes are approximately parallel to the layers of the lobula and the new Z axis is normal to the layers and goes through the retinotopic center of the lobula. This new coordinate system was obtained by performing principal component analysis on the postsynaptic terminals of the 4 LT1 neurons in the lobula. LT1 neurons have a dense, monostratified dendrite in the lobula layer 2 that covers the entire tangential extent of the lobula (Fischbach and Dittrich, 1989), which can be used as a landmark. We then calculated three standard deviations of the positions of presynapses of each lobula intrinsic terminals along each dimension of the new coordinate system, which respectively characterized the spatial spread of the terminals in the two tangential dimensions (PC1 and 2) and the normal dimension (PC3). Third, to identify the layer affiliation of each synaptic terminal, we first fit a surface model to the positions of the presynaptic terminals of LT1, which predicted PC3 position of each synapse with a bivariate quadratic formula of PC1 and PC2. The least square fit resulted in R^2^ = 0.74. Then, for each postsynapse location of the lobula intrinsic terminals, we calculated deviation between its actual PC3 position and the prediction from the quadratic model, which was interpreted as the relative depth of the synapse with respect to the layer 2 under the assumption that the layer boundaries of the lobula can be approximated as parallel quadric manifolds (positive deviations corresponding to deeper layers). For each lobula intrinsic terminal, we counted the numbers of the synapses whose fell in eleven 5 µm bins ranging from -10 µm to 45 µm. Finally, we ran a hierarchical agglomerative clustering on the 977 x 140 connectivity-morphology-innervation matrix and extracted 15 clusters, whose membership sizes varied from 18 to 148 cells. We then visualized the all neurons in each cluster on neuPrint explorer (Clements et al., 2020) (**Fig. S3C**), and examined their morphology while referencing anatomical literature to identify putative cell types (Fischbach and Dittrich, 1989). The code to run the clustering analysis can be found on our GitHub repository (https://github.com/ClarkLabCode/LPLC1ConnectomeAnalysis), and the complete list of the cells analyzed with their cluster affiliation is provided in **Supplementary File 2**. The list of visually annotated T2 and T3 cells can be found in **Supplementary File 3**.

### Connectomic identification and split Gal4 generation for downstream targets of LPLC1

Major downstream neuron types of LPLC1 were identified in the hemibrain v1.1 dataset (Scheffer et al., 2020) through the neuprint website (Clements et al., 2020). Since there were no preexisting selective Gal4 drives to label PLP219 and PVLP112/113, we created a new split Gal4 lines by screening for hemidrivers targeting these cell types using color depth maximum intensity projection search (Otsuna et al., 2018) running on multi-color flip out image library (Meissner et al., 2020) on the NeuronBridge website (Clements et al., 2020).

### Immunohistochemistry

The tissues were dissected out in PBS, fixed in 4% paraformaldehyde for 15 minutes, washed three times for 20 minutes, blocked with 5% normal goat serum for another 20 minutes, and incubated with primary antibodies (mouse anti-Brp, 1:25; chicken anti-GFP, 1:50) in PBST (PBS with 0.2% Triton-X) for 24 hours. After another 3 washes, the tissues were incubated with secondary antibodies (goat anti-mouse AF633, 1:250; goat anti-Chicken AF488, 1:250). 5% normal goat serum was also added to the primary and secondary antibody solutions. The tissues were then mounted on glass microscope slides with Vectashield mounting medium, and imaged with a Zeiss confocal microscope.

### Quantification and statistical analysis

For statistical purposes, each fly or cell was counted as an independent measurement, as noted in the figure captions. *p*-values presented are from Wilcoxon sign-rank tests (within-fly comparisons across stimulus conditions), rank-sum tests (across-fly comparisons across populations), Friedman test (within-fly comparisons across more than 3 stimulus conditions), or 2-way ANOVA (across-population comparison of tuning curves where only the existence of the population main effect matters). The tests are all as noted in the figure captions.

---

**Supplementary Figure 1.**
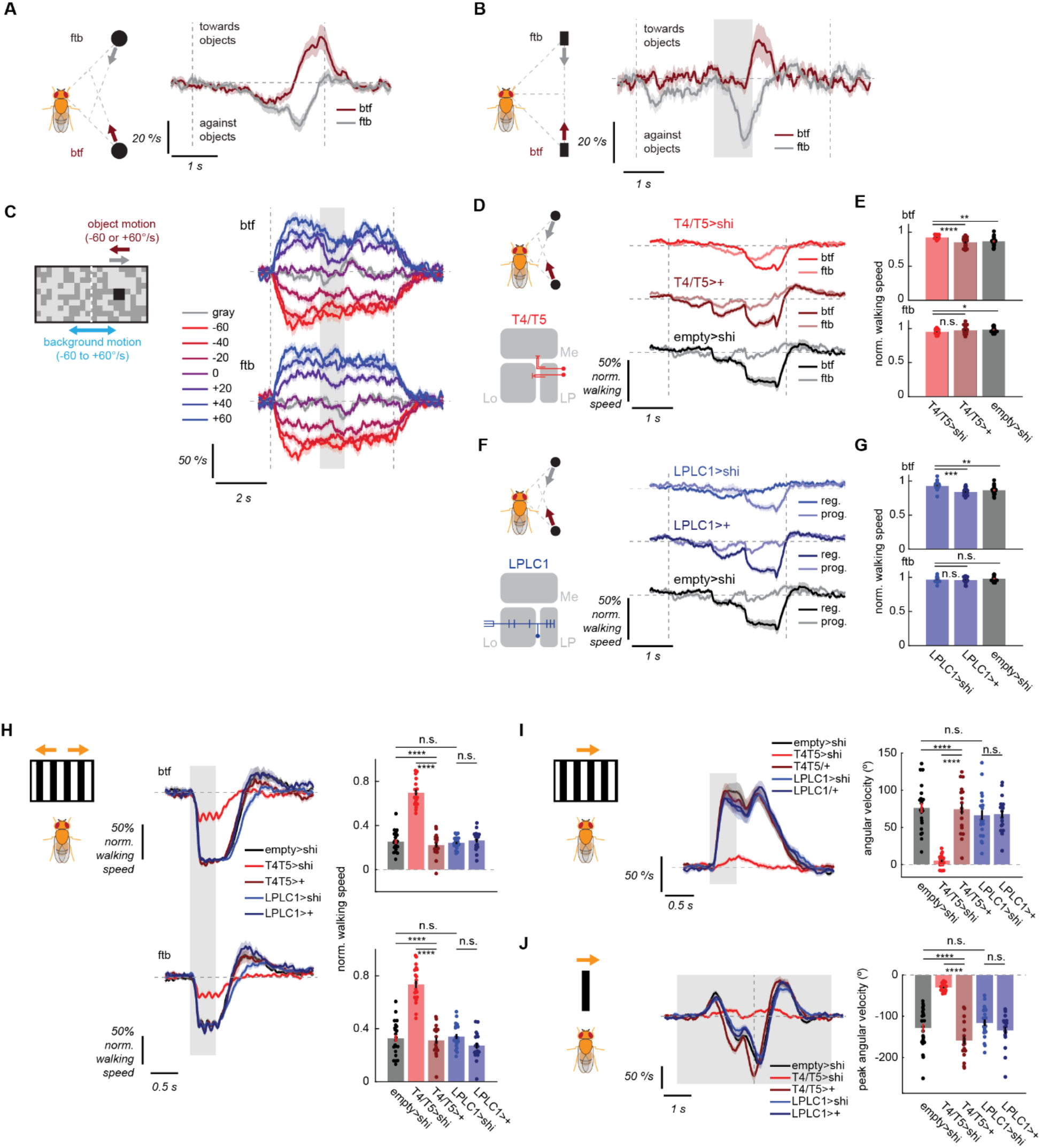
Additional behavioral characterization. (Related to Figs. 1, 2) (A-C) Turning responses of wildtype flies to the (A) approach stimuli, (B) parallel stimuli, and (C) squares paired with rotating backgrounds, by stimulus directions and background velocities. (D-G) Normalized walking responses of flies to the approach stimuli with (D, E) T4/T5 or (F, G) LPLC1 silenced and their respective genetic controls, either (D, F) over time or (E, G) time-averaged, as in **Fig. 1G, H**. (H-J) Examples of T4/T5-dependent behaviors where LPLC1 is dispensable. (H) Slowing responses of flies with T4/T5 or LPLC1 silencing and their controls to translational gratings either back-to-front or front-to-back, either (*left*) over time or (*right*) time-averaged. (I) Optomotor turning responses of flies with T4/T5 or LPLC1 silencing and their controls to drifting gratings, either (*left*) over time or (*right*) time-averaged. (J) Aversive turning responses of flies with T4/T5 or LPLC1 silencing and their controls to fast translating vertical bars, either (*left*) over time or (*right*) peak turning amplitudes. Error bars and shades around mean traces all indicate standard error of the mean. (A) N = 21 flies. (B) N = 19 flies. (C) N = 19 flies. (D, E) N = 20 (T4/T5>shi), 23 (T4T5>+), 21 (empty>shi) flies. (C, D) N = 16 (LPLC1>shi), 21 (LPLC1>+), 21 (empty>shi) flies. (H-J) N = 19 (T4/T5>shi), 17 (T4/T5>+), 20 (LPLC1>shi), 17 (LPLC1>+), 22 (empty>shi) flies. n. s.: not significant (p > .05); *: p < .05; **: p < .01; ***: p < .001; ****: p < .0001 in Wilcoxon rank sum test.

**Supplementary Figure 2.**
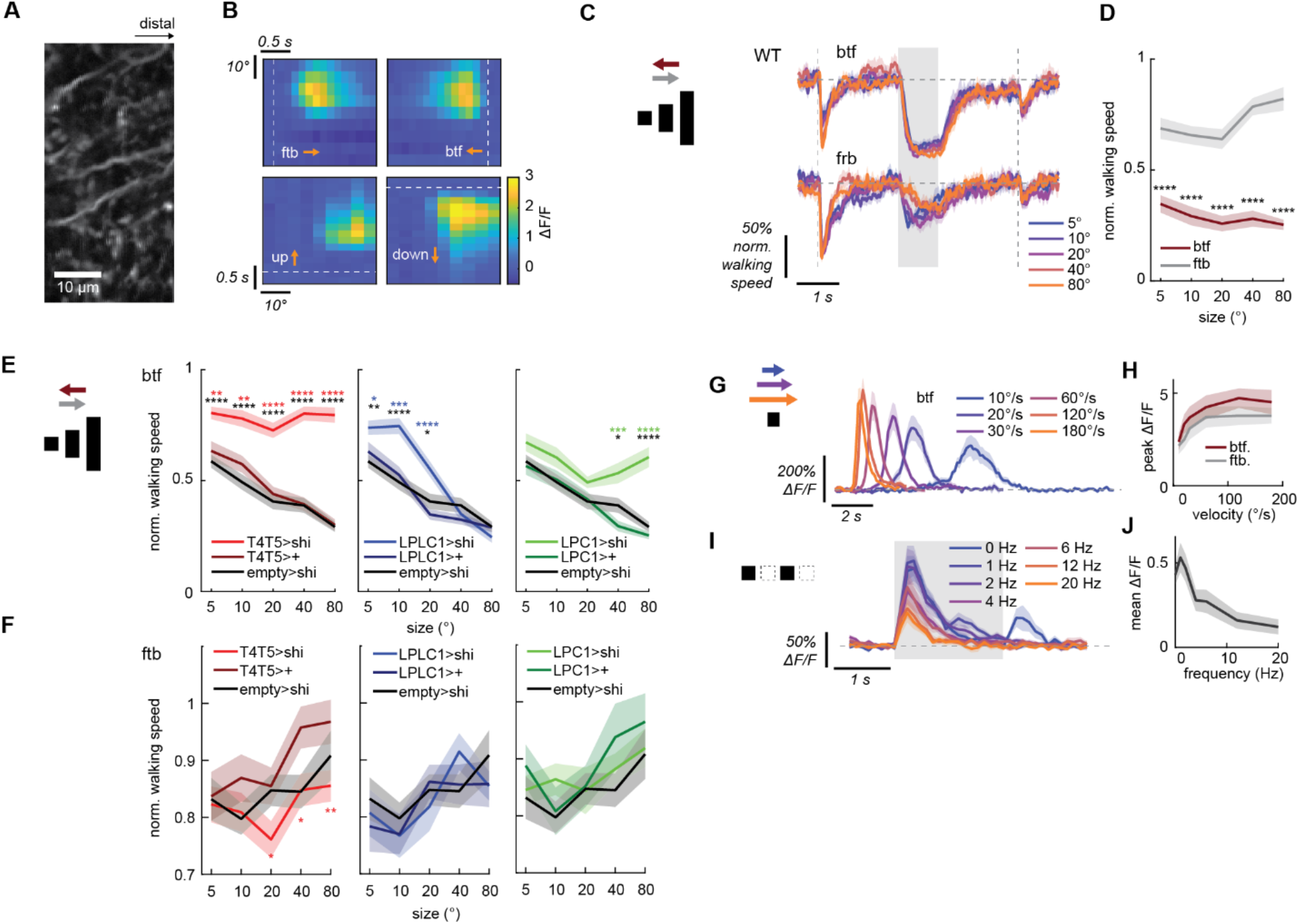
Additional characterizations of LPLC1 cell response properties and their behavioral consequences. (Related to Fig. 3.) (A) A representative time-averaged image of LPLC1 lobula neurites during single cell recordings. (B) Responses of an example cell to receptive field mapping stimuli (a 10° x 10° translating black square). The square swept a 40° x 40° area around the approximated RF center in the four directions with the resolution of 5°. (C, D) Wildtype fly slowing response to horizontally translating objects with different heights in either direction, either (C) over time or (D) as functions of the heights. Flies slow more in response to objects moving back-to-front across the all sizes tested. (E) Slowing responses of flies with (*left*) T4/T5, (*middle*) LPLC1, or (*right*) LPC1 silenced and their respective controls to objects with various heights, moving back-to-front. While silencing of T4/T5 reduces slowing across the all sizes tested, LPLC1 and LPC1 silencing only affects slowing caused by short and tall objects, respectively, revealing complementary contributions to behavior of these two visual projection neurons. (F) The same as (E), but for objects moving front-to-back, where these manipulations had little effect. (G-J) Calcium responses of LPLC1 neurons to small squares either (G, H) translating at various velocities or (I, J) flickering on the spot at various temporal frequencies, either over time or as functions of the velocity/temporal frequency. Error bars and shading around mean traces all indicate standard error of the mean across (C-F) flies or (G-J) cells. (C, D) N = 16 flies. (E, F) N = 21 (T4/T5>shi), 18 (T4/T5>+), 19 (LPLC1>shi), 18 (LPLC1>+), 20 (LPC1>shi), 19 (LPC1>+), 21 (empty>shi) flies. (G, H) N = 11 cells. (I, J) N = 12 cells. n. s.: not significant (p > .05); *: p < .05; **: p < .01; ***: p < .001; ****: p < .0001 in Wilcoxon signed-rank (D) or rank sum test (E, F).

**Supplementary Figure 3.**
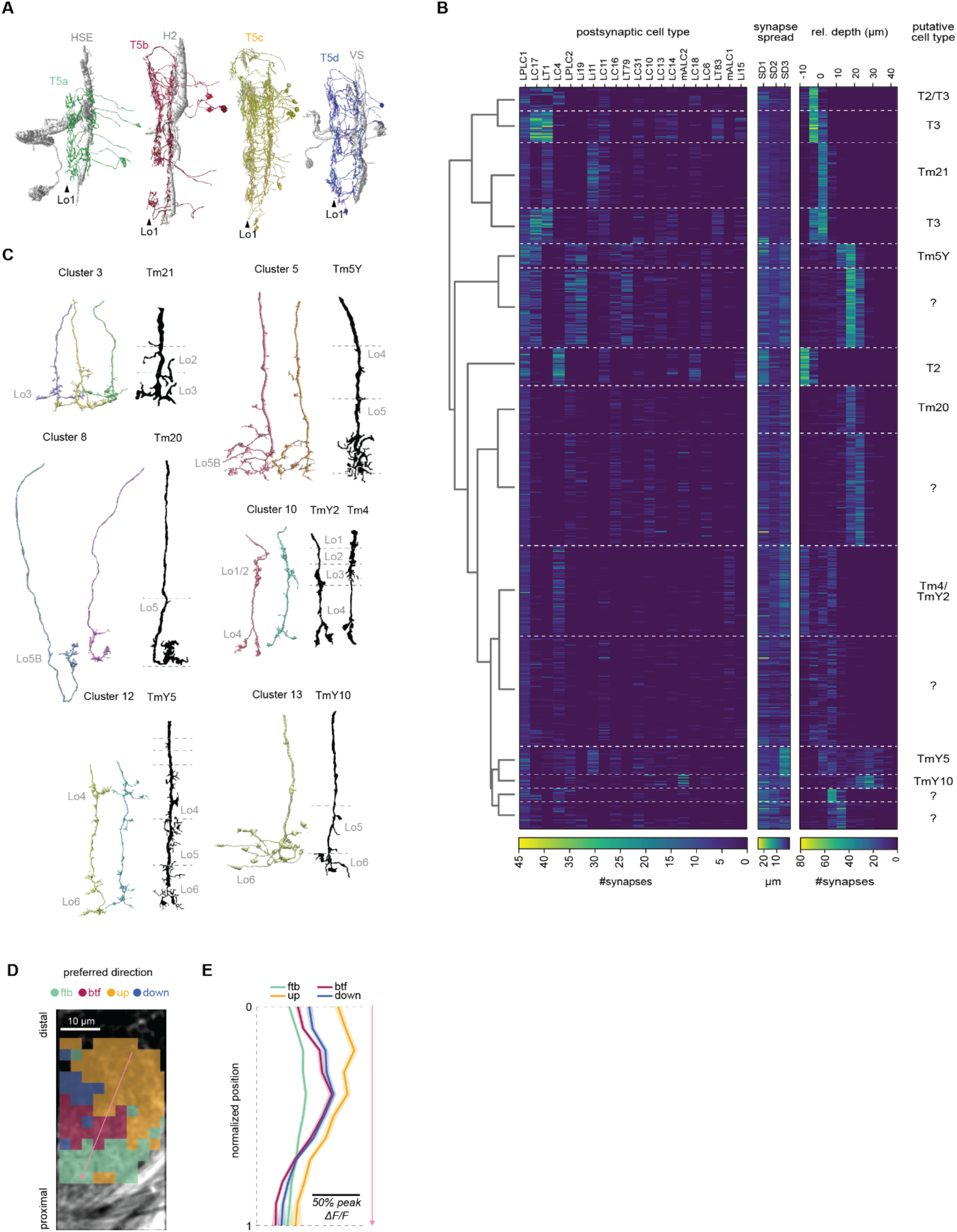
Additional characterization of LPLC1 inputs. (Related to Fig. 4) (A) Example T5 cells grouped by their subtypes, with their postsynaptic tangential neurons. (B) Three feature matrices used for lobula terminal clustering, representing (*left*) connectivity to downstream cell types, (*middle*) spatial spread of synapses along the three axes, and (*right*) innervation depth relative to LT1 are shown. Each row corresponds to a single terminal fragment. The dendrogram on the left shows the result of the agglomerative clustering. The putative cell type labels are shown on the right. Only the top 20 postsynaptic cell types are visualized. See also **Supplementary File 2**. (C) Reconstructed single-cell morphology of example neurons from clusters that resembled known cell types, alongside previously published Golgi-staining (in black) (Fischbach and Dittrich, 1989). (D) An example image of lobula plate expressing iGluSnFR panneuronally (nSyb > iGluSnFR), in which ROIs are color coded according to the direction of the bar to which they responded best, similar to Fig. 4I. (E) Peak glutamatergic signals in lobula plate, as functions of normalized positions of ROIs along the layers of lobula plate, measured from the distal most layer. Similar to Fig. 4J. Glutamate signals in the near layer 1 (normalized position 1) responded more to front-to-back than back to front, while closer to layer 2, the back-to-front signals are larger, suggesting intra-layer glutamate signaling. N = 17 (flies), 1595 (ROIs).

**Supplementary Figure 4.**
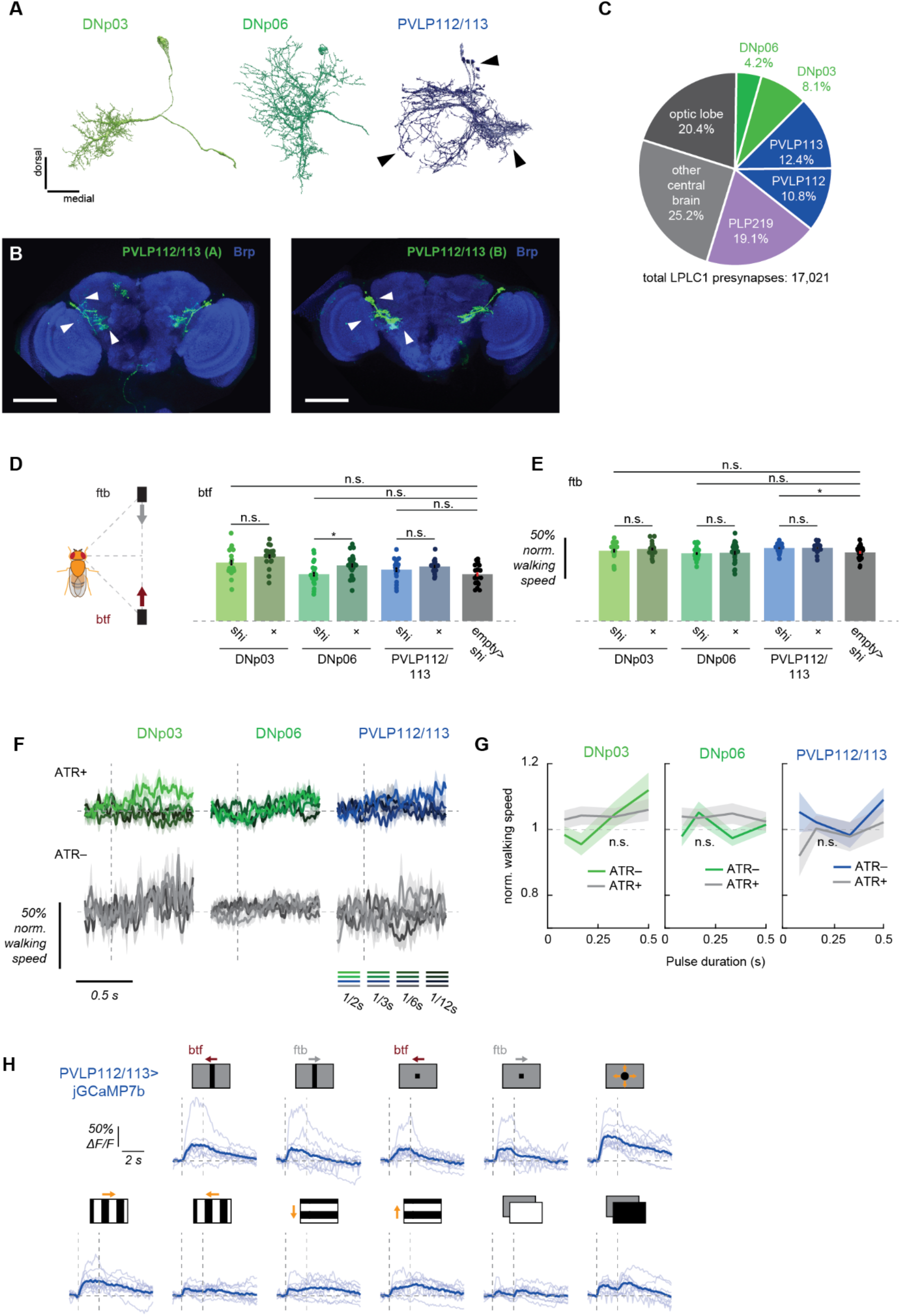
Neurons downstream of LPLC1. (Related to Fig. 5) (A) Reconstructed morphology of DNp03, DNp06, and PLP112/113 from the hemibrain dataset (Scheffer et al., 2020). (B) The expression patterns of two newly generated split Gal4 lines to label PVLP112/113 neurons, visualized with UAS-myr::GFP. (*left*) R72A10AD; VT002042DBD (*right*) R72A10AD; VT019749DBD. The first driver was used for the behavioral experiments. (C) Counts of LPLC1 output synapses by postsynaptic cell types. (D, E) Time-averaged normalized walking responses of flies to the parallel stimulus in (D) back-to-front and (E) front-to-back directions, with noted downstream neuron silenced and corresponding controls. (F, G) Normalized walking responses of flies expressing Chrimson in PVLP112/113, DNp03, or DNp06 to pulses of green lights, visualized (F) over time or (G) averaged over time. (H) Individual (light blue) and fly-averaged (dark blue) calcium responses of PVLP112/113 population over time to a variety of visual stimuli (horizontally moving bars and squares, looming, square wave gratings, full-field flashes). Leftward in the stimulus schematics corresponds to the back-to-front direction. Error bars and shades around mean traces all indicate standard error of the mean. (D, E) N = 17 (DNp03>shi), 19 (DNp03>+), 19 (DNp06>shi), 24 (DNp06>+), 19 (PVLP112/113>shi), 17 (PVLP112/113>+), 22 (empty>shi). (F, G) N = 14 (DNp03, ATR+), 13 (DNp03, ATR-), 13 (DNp06, ATR+), 15 (DNp03, ATR-), 12 (PVLP112/113, ATR+), 6 (PVLP112/113, ATR-). (H) N = 10. n. s.: not significant (p >.05); *: p < .05; **: p < .01; ***: p < .001; ****: p < .0001 in Wilcoxon rank sum test (D, E) and 2-way analysis of variance (ANOVA) (G; the main effect of ATR conditions).

**Supplementary Table 1.**
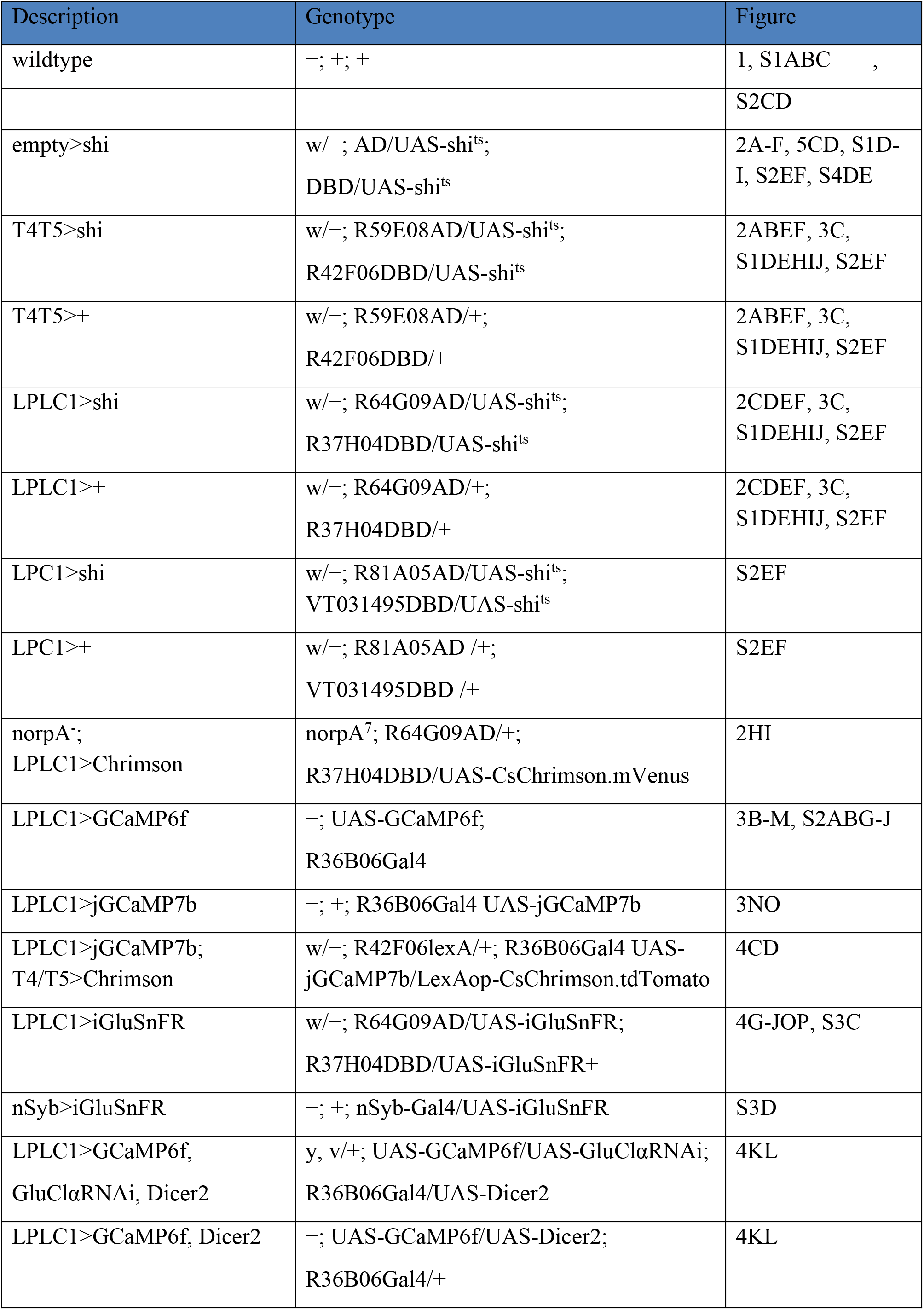
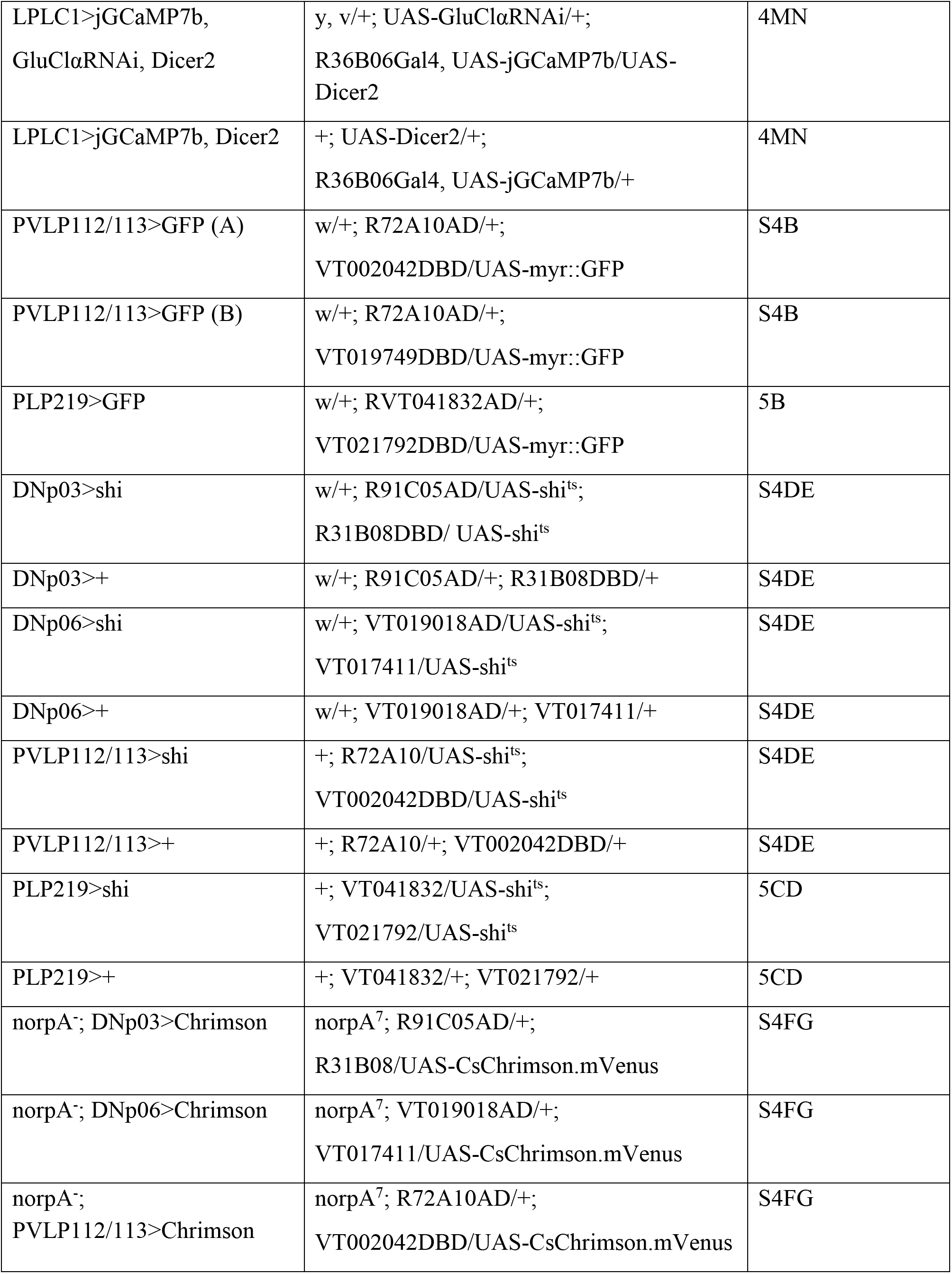
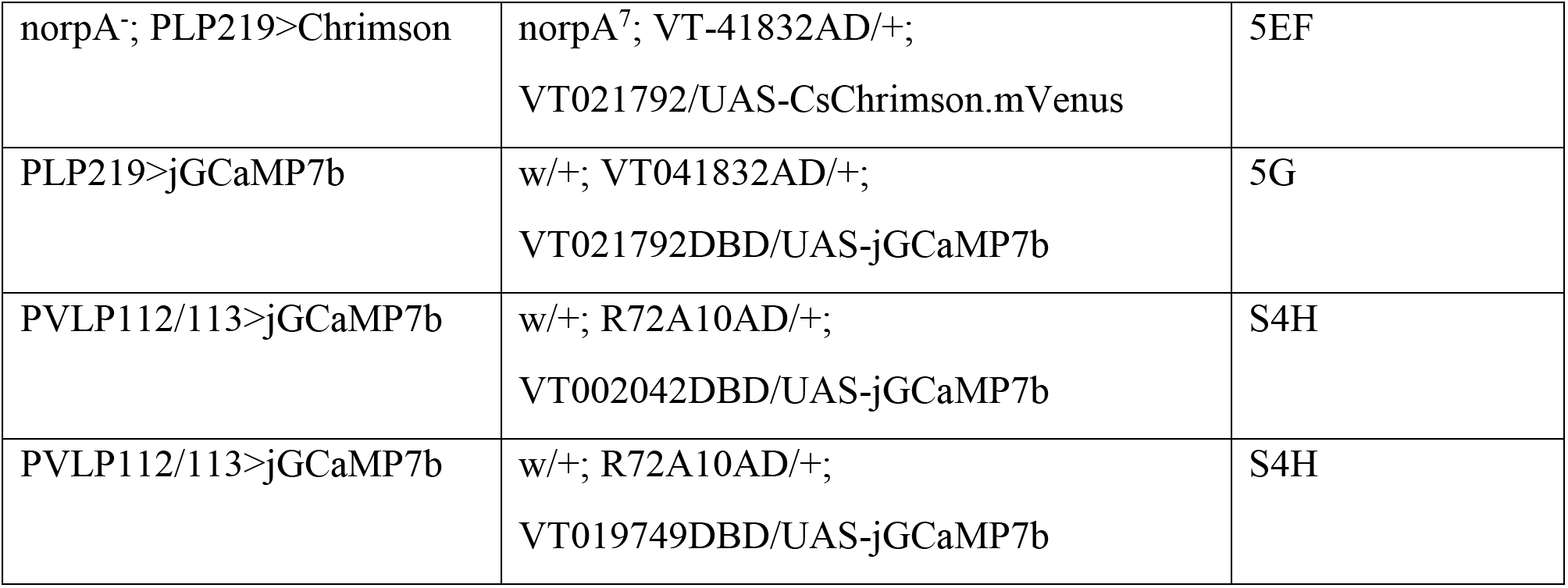
Genotypes of flies used in the experiments.

| Description | Genotype | Figure |
| --- | --- | --- |
| wildtype | +; +; + | 1, S1ABC , |
|  |  | S2CD |
| empty>shi | w/+; AD/UAS-shi<sup>ts</sup>;<br>DBD/UAS-shi<sup>ts</sup> | 2A-F, 5CD, S1D-I, S2EF, S4DE |
| T4T5>shi | w/+; R59E08AD/UAS-shi<sup>ts</sup>;<br>R42F06DBD/UAS-shi<sup>ts</sup> | 2ABEF, 3C, S1DEHIJ, S2EF |
| T4T5>+ | w/+; R59E08AD/+;<br>R42F06DBD/+ | 2ABEF, 3C, S1DEHIJ, S2EF |
| LPLC1>shi | w/+; R64G09AD/UAS-shi<sup>ts</sup>;<br>R37H04DBD/UAS-shi<sup>ts</sup> | 2CDEF, 3C, S1DEHIJ, S2EF |
| LPLC1>+ | w/+; R64G09AD/+;<br>R37H04DBD/+ | 2CDEF, 3C, S1DEHIJ, S2EF |
| LPC1>shi | w/+; R81A05AD/UAS-shi<sup>ts</sup>;<br>VT031495DBD/UAS-shi<sup>ts</sup> | S2EF |
| LPC1>+ | w/+; R81A05AD /+;<br>VT031495DBD /+ | S2EF |
| norpA<sup>-</sup>;<br>LPLC1>Chrimson | norpA<sup>7</sup>; R64G09AD/+;<br>R37H04DBD/UAS-CsChrimson.mVenus | 2HI |
| LPLC1>GCaMP6f | +; UAS-GCaMP6f;<br>R36B06Gal4 | 3B-M, S2ABG-J |
| LPLC1>jGCaMP7b | +; +; R36B06Gal4 UAS-jGCaMP7b | 3NO |
| LPLC1>jGCaMP7b;<br>T4/T5>Chrimson | w/+; R42F06lexA/+; R36B06Gal4 UAS-jGCaMP7b/LexAop-CsChrimson.tdTomato | 4CD |
| LPLC1>iGluSnFR | w/+; R64G09AD/UAS-iGluSnFR;<br>R37H04DBD/UAS-iGluSnFR+ | 4G-JOP, S3C |
| nSyb>iGluSnFR | +; +; nSyb-Gal4/UAS-iGluSnFR | S3D |
| LPLC1>GCaMP6f,<br>GluClαRNAi, Dicer2 | y, v/+; UAS-GCaMP6f/UAS-GluClαRNAi;<br>R36B06Gal4/UAS-Dicer2 | 4KL |
| LPLC1>GCaMP6f, Dicer2 | +; UAS-GCaMP6f/UAS-Dicer2;<br>R36B06Gal4/+ | 4KL |
| LPLC1>jGCaMP7b,<br>GluClαRNAi, Dicer2 | y, v/+; UAS-GluClαRNAi/+;<br>R36B06Gal4, UAS-jGCaMP7b/UAS-Dicer2 | 4MN |
| LPLC1>jGCaMP7b, Dicer2 | +; UAS-Dicer2/+;<br>R36B06Gal4, UAS-jGCaMP7b/+ | 4MN |
| PVLP112/113>GFP (A) | w/+; R72A10AD/+;<br>VT002042DBD/UAS-myr::GFP | S4B |
| PVLP112/113>GFP (B) | w/+; R72A10AD/+;<br>VT019749DBD/UAS-myr::GFP | S4B |
| PLP219>GFP | w/+; RVT041832AD/+;<br>VT021792DBD/UAS-myr::GFP | 5B |
| DNp03>shi | w/+; R91C05AD/UAS-shi<sup>ts</sup>;<br>R31B08DBD/ UAS-shi<sup>ts</sup> | S4DE |
| DNp03>+ | w/+; R91C05AD/+; R31B08DBD/+ | S4DE |
| DNp06>shi | w/+; VT019018AD/UAS-shi<sup>ts</sup>;<br>VT017411/UAS-shi<sup>ts</sup> | S4DE |
| DNp06>+ | w/+; VT019018AD/+; VT017411/+ | S4DE |
| PVLP112/113>shi | +; R72A10/UAS-shi<sup>ts</sup>;<br>VT002042DBD/UAS-shi<sup>ts</sup> | S4DE |
| PVLP112/113>+ | +; R72A10/+; VT002042DBD/+ | S4DE |
| PLP219>shi | +; VT041832/UAS-shi<sup>ts</sup>;<br>VT021792/UAS-shi<sup>ts</sup> | 5CD |
| PLP219>+ | +; VT041832/+; VT021792/+ | 5CD |
| norpA<sup>-</sup>; DNp03>Chrimson | norpA<sup>7</sup>; R91C05AD/+;<br>R31B08/UAS-CsChrimson.mVenus | S4FG |
| norpA<sup>-</sup>; DNp06>Chrimson | norpA<sup>7</sup>; VT019018AD/+;<br>VT017411/UAS-CsChrimson.mVenus | S4FG |
| norpA<sup>-</sup>;<br>PVLP112/113>Chrimson | norpA<sup>7</sup>; R72A10AD/+;<br>VT002042DBD/UAS-CsChrimson.mVenus | S4FG |
| norpA<sup>-</sup>; PLP219>Chrimson | norpA<sup>7</sup>; VT-41832AD/+;<br>VT021792/UAS-CsChrimson.mVenus | 5EF |
| PLP219>jGCaMP7b | w/+; VT041832AD/+;<br>VT021792DBD/UAS-jGCaMP7b | 5G |
| PVLP112/113>jGCaMP7b | w/+; R72A10AD/+;<br>VT002042DBD/UAS-jGCaMP7b | S4H |
| PVLP112/113>jGCaMP7b | w/+; R72A10AD/+;<br>VT019749DBD/UAS-jGCaMP7b | S4H |

**Supplementary Table 2.**
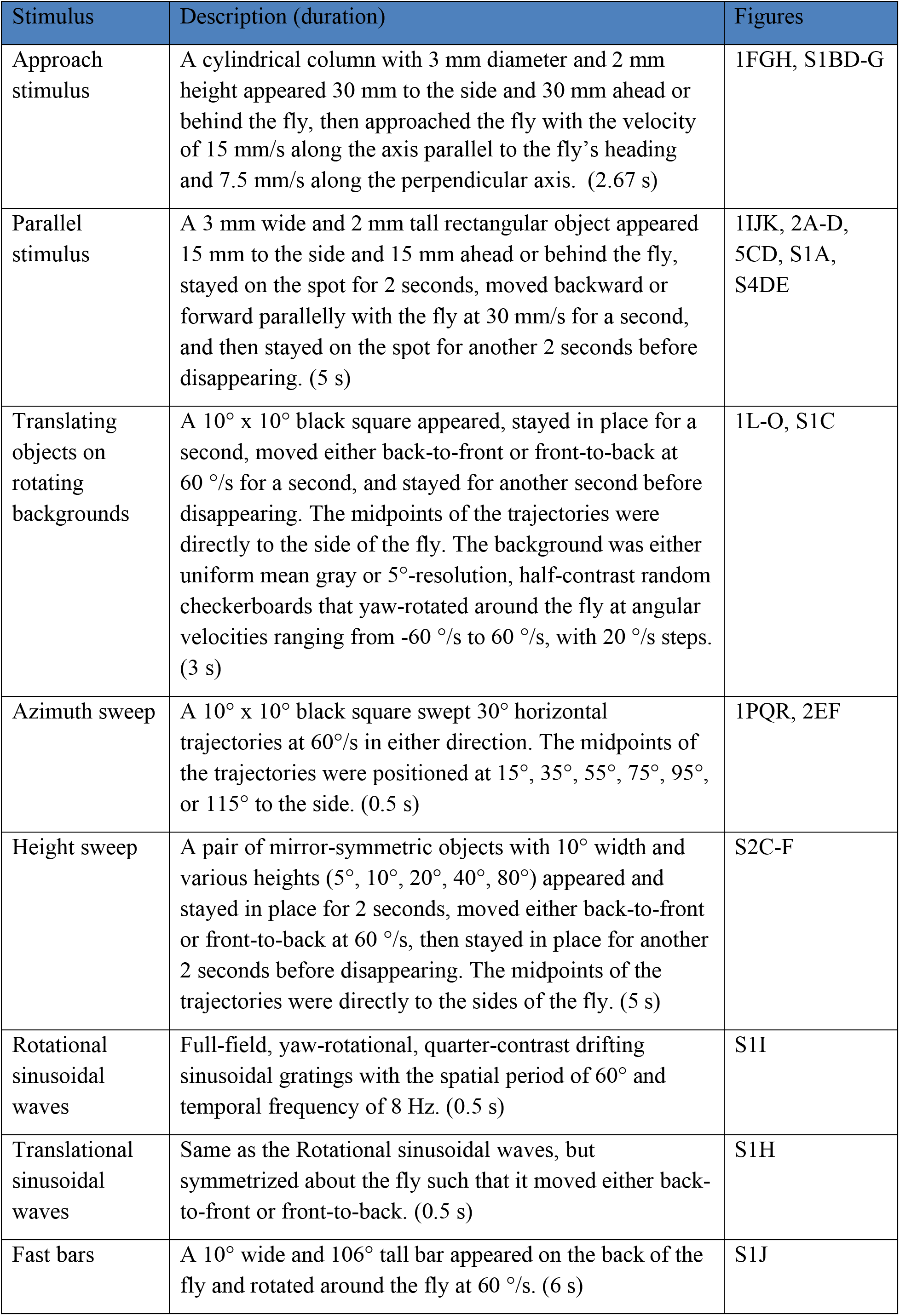
Stimuli used in the behavioral experiments.

**Supplementary Table 3.**
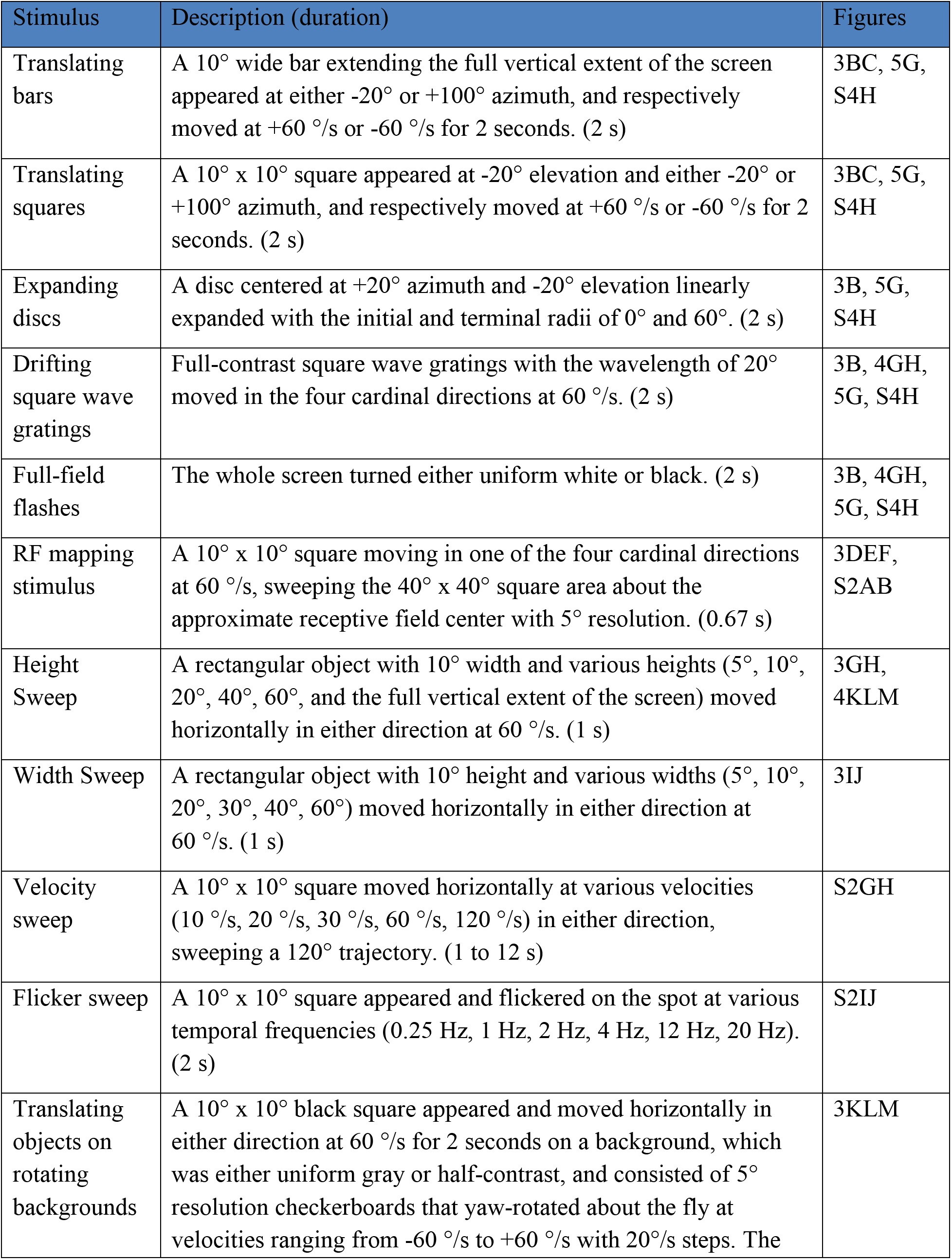
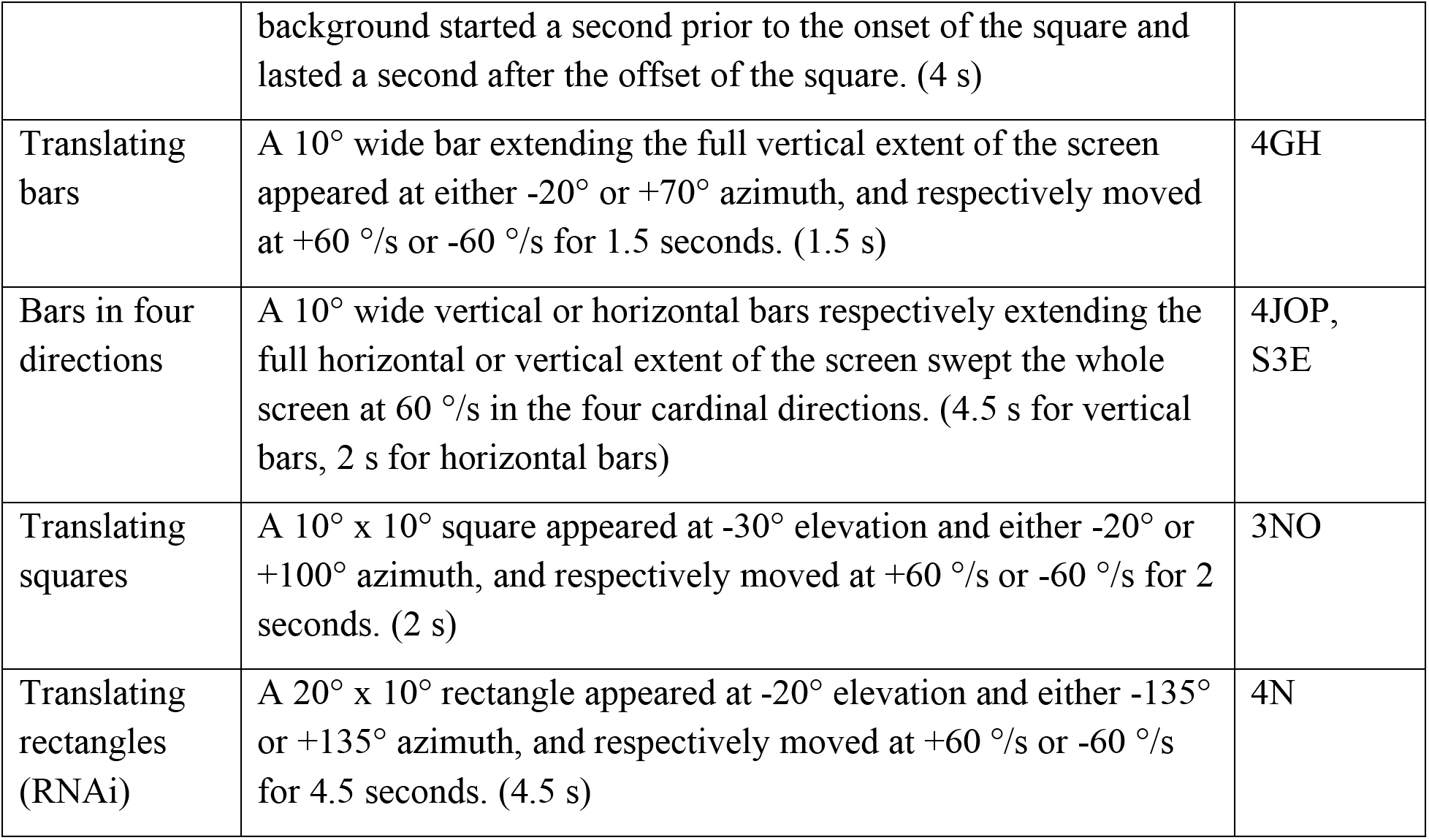
Stimuli used in the imaging experiments.

| Stimulus | Description (duration) | Figures |
| --- | --- | --- |
| Translating bars | A 10° wide bar extending the full vertical extent of the screen appeared at either -20° or +100° azimuth, and respectively moved at +60 °/s or -60 °/s for 2 seconds. (2 s) | 3BC, 5G, S4H |
| Translating squares | A 10° x 10° square appeared at -20° elevation and either -20° or +100° azimuth, and respectively moved at +60 °/s or -60 °/s for 2 seconds. (2 s) | 3BC, 5G, S4H |
| Expanding discs | A disc centered at +20° azimuth and -20° elevation linearly expanded with the initial and terminal radii of 0° and 60°. (2 s) | 3B, 5G, S4H |
| Drifting square wave gratings | Full-contrast square wave gratings with the wavelength of 20° moved in the four cardinal directions at 60 °/s. (2 s) | 3B, 4GH, 5G, S4H |
| Full-field flashes | The whole screen turned either uniform white or black. (2 s) | 3B, 4GH, 5G, S4H |
| RF mapping stimulus | A 10° x 10° square moving in one of the four cardinal directions at 60 °/s, sweeping the 40° x 40° square area about the approximate receptive field center with 5° resolution. (0.67 s) | 3DEF, S2AB |
| Height Sweep | A rectangular object with 10° width and various heights (5°, 10°, 20°, 40°, 60°, and the full vertical extent of the screen) moved horizontally in either direction at 60 °/s. (1 s) | 3GH, 4KLM |
| Width Sweep | A rectangular object with 10° height and various widths (5°, 10°, 20°, 30°, 40°, 60°) moved horizontally in either direction at 60 °/s. (1 s) | 3IJ |
| Velocity sweep | A 10° x 10° square moved horizontally at various velocities (10 °/s, 20 °/s, 30 °/s, 60 °/s, 120 °/s) in either direction, sweeping a 120° trajectory. (1 to 12 s) | S2GH |
| Flicker sweep | A 10° x 10° square appeared and flickered on the spot at various temporal frequencies (0.25 Hz, 1 Hz, 2 Hz, 4 Hz, 12 Hz, 20 Hz). (2 s) | S2IJ |
| Translating objects on rotating backgrounds | A 10° x 10° black square appeared and moved horizontally in either direction at 60 °/s for 2 seconds on a background, which was either uniform gray or half-contrast, and consisted of 5° resolution checkerboards that yaw-rotated about the fly at velocities ranging from -60 °/s to +60 °/s with 20°/s steps. The | 3KLM |
|  | background started a second prior to the onset of the square and lasted a second after the offset of the square. (4 s) |  |
| Translating bars | A 10° wide bar extending the full vertical extent of the screen appeared at either -20° or +70° azimuth, and respectively moved at +60 °/s or -60 °/s for 1.5 seconds. (1.5 s) | 4GH |
| Bars in four directions | A 10° wide vertical or horizontal bars respectively extending the full horizontal or vertical extent of the screen swept the whole screen at 60 °/s in the four cardinal directions. (4.5 s for vertical bars, 2 s for horizontal bars) | 4JOP, S3E |
| Translating squares | A 10° x 10° square appeared at -30° elevation and either -20° or +100° azimuth, and respectively moved at +60 °/s or -60 °/s for 2 seconds. (2 s) | 3NO |
| Translating rectangles (RNAi) | A 20° x 10° rectangle appeared at -20° elevation and either -135° or +135° azimuth, and respectively moved at +60 °/s or -60 °/s for 4.5 seconds. (4.5 s) | 4N |

**Supplementary Table 4.** Probe stimuli used in the imaging experiments.

| Stimulus | Description (duration) | Figures |
| --- | --- | --- |
| Vertical bars | A 10° wide bar extending the full vertical extent of the screen appeared at either -20° or +70° azimuth, and respectively moved at +60 °/s or -60 °/s for 1.5 seconds. (1.5 s) | 4GH |
| Bars in four directions | A 10° wide vertical or horizontal bars respectively extending the full horizontal or vertical extent of the screen swept the whole screen at 60 °/s in the four cardinal directions. (4.5 s for vertical bars, 2 s for horizontal bars) | 4JOP, S3E |
| Translating squares | A 10° x 10° square appeared at -30° elevation and either -20° or +100° azimuth, and respectively moved at +60 °/s or -60 °/s for 2 seconds. (2 s) | 3NO |
| Translating rectangles (RNAi) | A 20° x 10° rectangle appeared at -20° elevation and either -135° or +135° azimuth, and respectively moved at +60 °/s or -60 °/s for 4.5 seconds. (4.5 s) | 4N |

